# Rapid catecholamine trafficking regulates neutrophil functions and neutrophil-platelet interactions

**DOI:** 10.1101/2025.01.27.634021

**Authors:** Jennifer Mohr, Anne Schmitz, Meshkat Dinarvand, Sangeetha Shankar, Bjoern F. Hill, Franziska Wulfert, Elsa Neubert, Magdalena Shumanska, Sofia Kaushik, Linda Kartaschew, Ivan Bogeski, James Daniel, Sebastian Jung, Johannes Eble, Luise Erpenbeck, Sebastian Kruss

## Abstract

Neutrophils are key effector cells of the innate immune system that respond to small signaling molecules regulating immune responses. For a long time, similarities between neuronal and immune cells have been discussed. Here, we show that human neutrophils rapidly take up, package and use catecholamine neurotransmitters such as dopamine or epinephrine via the machinery known from neurons. Uptake and release of catecholamines as well as trafficking and packaging into MPO/VMAT2-positive primary vesicles is visualized with false fluorescent neurotransmitters. We also directly image the fast (> 10 s) and transient release of catecholamines from neutrophils with near infrared fluorescent nanosensors. Serotonin or activated platelets trigger calcium (Ca^2+^) signaling and consequently exocytosis of catecholamines. They reduce NET-formation but increase platelet aggregation. Thus, we establish similarities between neurons and neutrophils and identify a paracrine neutrophil-platelet feedback loop relevant for inflammatory and coagulatory conditions.

## Introduction

Neutrophilic granulocytes (neutrophils) are key effector cells of the innate immune system and possess multiple highly effective immune-defense mechanisms such as formation of neutrophil extracellular traps (NETs).(*1, 2*) Dysregulated or excessive neutrophil activation and NETosis is implicated in a plethora of inflammatory and malignant diseases inducing extensive acute tissue damage as well as thromboembolic events.(*1, 3*) NET formation can be initiated by activated platelets, and neutrophils together with activated platelets participate in thromboinflammation(*3*), which can cause severe tissue damage in the host.(*4*) Therefore, neutrophil activation and particularly NET formation must be tightly regulated, both on a global, systemic level but also during transient cell-cell interactions. Communication via small signaling molecules that diffuse rapidly between cells contributes to these interactions. Catecholamine (CA) neurotransmitters like dopamine (DA), epinephrine (E), or norepinephrine (NE) have long been known as neuromodulators in the nervous and endocrine system.(*5*) During exocytosis, secretory vesicles fuse with the cell membrane, which releases the vesicular content into the extracellular space.(*6*) Exocytosis is important for various biological processes such as neurotransmitter release or hormone secretion.(*7*) It requires vesicular docking, priming, and fusion, which is performed by a complex machinery of proteins such as soluble N-ethylmaleimide-sensitive factor attachment receptors (SNAREs).(*8*) In immune cells such as neutrophils, exocytosis is associated with the release of antimicrobial substances and enzymes to combat infections.(*9*)

There is growing evidence that CAs are also involved in the peripheral immune system. For example, follicular T helper cells produce and release DA, which is recognized by B cells in germinal centers and accelerates productive synapse formation.(*10*) Macrophages, mast cells, and lymphocytes either produce DA or possess DA receptors.(*11*–*14*) Adrenergic signaling has been reported in granulocytes, macrophages, dendritic cells, natural killer cells and lymphocytes. (*15, 16*) For neutrophils, there is conflicting information about the expression of DA receptors while the expression of adrenergic receptors has previously been shown.(*14, 17*) The role of CAs in neutrophils remains enigmatic. It has been described that neutrophils can synthesize and release CAs upon lipopolysaccharide (LPS) challenge and that this may regulate inflammatory responses via adrenoreceptors.(*18, 19*) However, the mechanism of CA uptake, packaging, storage, and release in analogy to neuronal cells has not been examined. One reason is that fast imaging of transient local changes in CA concentration is not possible with conventional biochemical methods.

CA can be detected by electrochemical approaches, mass spectrometry (MS), or high-pressure liquid chromatography (HPLC) but without the necessary spatial information. Genetically encoded sensors for DA have been used for brain studies but require transfection, which is not available for short-living human cells such as neutrophils.(*20*) An indirect method of studying the dynamics of CA is the use of fluorescent false neurotransmitters (FFNs). These fluorescent molecules have a similar structure to CAs and therefore high affinity for dopamine transporter (DAT) and vesicular monoamine transporter (VMAT).(*21*) Fluorescent nanosensors based on single-walled carbon nanotubes (SWCNTs) have shown great potential to image biochemical signaling.(*22*) SWCNTs consist of rolled-up layers of carbon and they fluoresce in the beneficial near-infrared (nIR) tissue transparency window.(*23*) They can be chemically tailored to recognize biomolecules such as neurotransmitters,(*24*) lipids, reactive oxygen species(*25*) or proteins(*26*) down to the single molecule level. These nanosensors have been employed to visualize single DA release events from pheochromocytoma (PC-12) cells,(*27*) primary dopaminergic neurons,(*28*) and exocytosis of serotonin (5-HT) from activated platelets.(*29*) Here, we show that neutrophils possess the CA machinery similar to dopaminergic neurons for uptake, packaging, storage, and the dynamics of CA exocytosis. We identify 5-HT as a trigger for CA exocytosis, image transient CA release by using nanosensors and show the impact of CA on neutrophil functions. These results provide evidence for a hitherto unrecognized, CA-based feedback loop between platelets and neutrophils.

## Results

### Neutrophils synthesize, store, and release catecholamines

First, we checked the presence of the catecholaminergic machinery known from neurons in neutrophils. To this end, human neutrophils were stained for VMAT2, DAT, and tyrosine hydroxylase (TH) (Fig. 1A). DAT transports CAs into neurons and VMAT2 is known to package CAs into vesicles(*30, 31*) and both are expressed in primary dopaminergic (mouse) neurons (Fig. S1). VMAT2 colocalized with myeloperoxidase (MPO), which indicates that CAs are stored in azurophilic granules. TH is the rate-limiting enzyme in CA synthesis and is used as a standard marker for dopaminergic neurons (Fig. 1A, Fig. S1). Further immunofluorescence staining as well as PCR was performed to identify CA processing and degrading enzymes (monoamine oxidase A, MAOA and monoamine oxidase B, MAOB) and catechol-O-methyltransferase (COMT, Fig. 1A, Fig. S2). The presence of the CA machinery does not imply that CAs are present in the cells. Additionally, given the presence of degrading enzymes (Fig. 1C), we wanted to check for CA degradation products. Therefore, an ELISA assay was performed to measure the intracellular concentration of CAs and their metabolites homovanillic acid (HVA) and vanillylmandelic acid (VMA) (Fig. 1B). CA in different concentrations as well as these metabolites were found in neutrophils. The number of, for example around 10^7^ DA molecules per cell suggests that approximately 300 vesicles could be filled if one assumes a quantal size of 33,000 DA molecules per cell as reported for neuronal vesicles.(*32*)

**Fig. 1.**
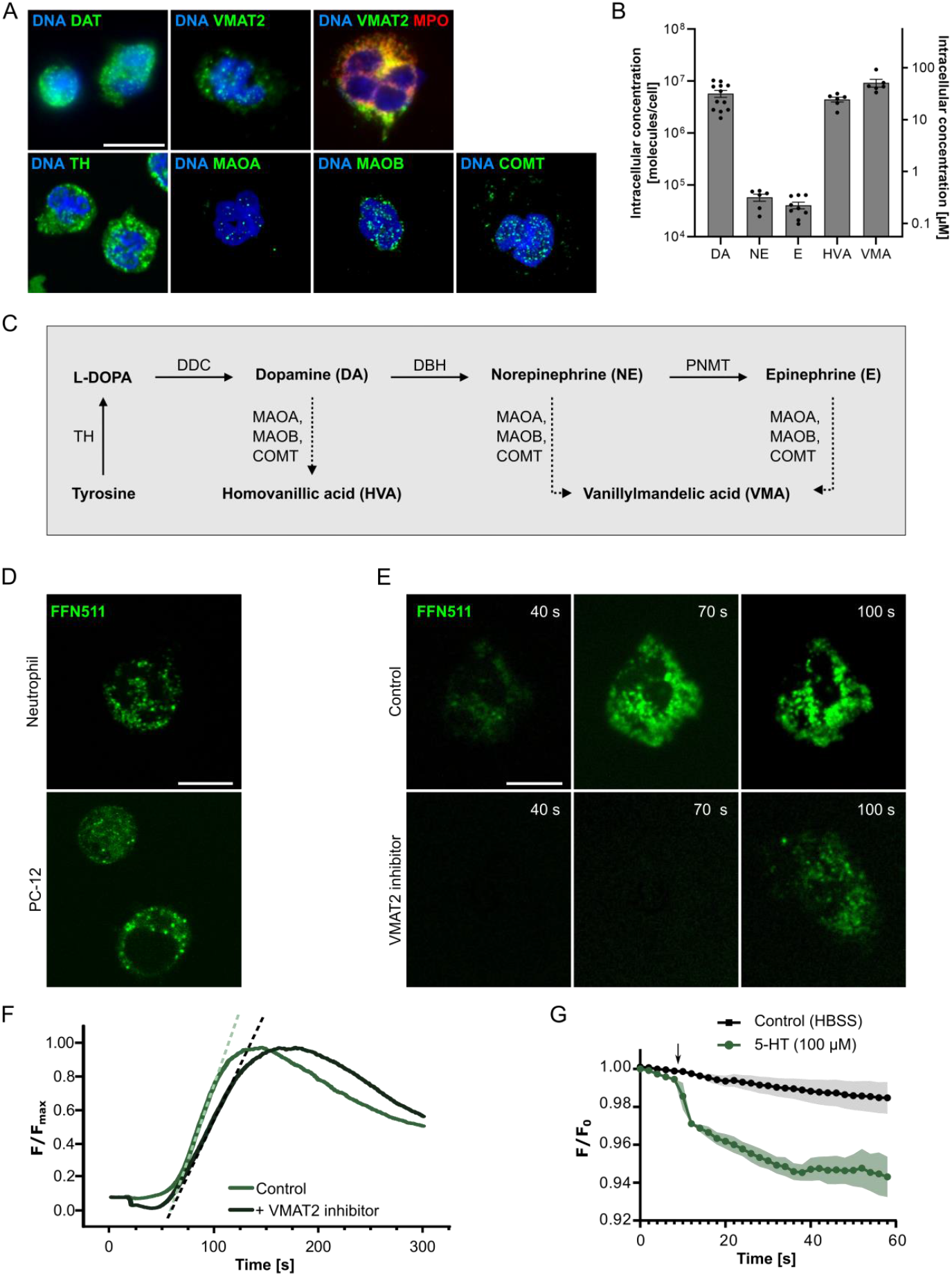
Neutrophils have the machinery to synthesize, degrade, take up, package, and release catecholamines. (**A**) Neutrophils are stained for dopamine transporter (DAT), vesicular monoamine transporter 2 (VMAT2), tyrosine hydroxylase (TH), monoamine oxidase A (MAOA), monoamine oxidase B (MAOB), catechol-O-methyltransferase (COMT) (all green), myeloperoxidase (MPO) (red), and DNA (blue). Scale bar is 10 μm. (**B**) Intracellular content of CAs and the metabolic products homovanillic acid (HVA) and vanillylmandelic acid (VMA) determined by an ELISA. (DA: n = 12, NE: n = 6, E: n = 9, mean ± SEM). Assumption for the neutrophil volume = (10 μm)^3^. (**C**) Scheme of catecholamine metabolism (DDC: DOPA decarboxylase, DBH: dopamine beta-hydroxylase, PNMT: phenylethanolamine N-methyltransferase). (**D**) Image of false fluorescent neurotransmitters (FFN511)-filled vesicles in live neutrophils and PC-12 cells (positive control). Scale bar is 10 μm. (**E**) FFN511 (2.5 nM) uptake of neutrophils with VMAT2 inhibition by tetrabenazine at different time points (**F**) Corresponding single-cell kinetic analysis (fluorescence intensity inside the cell) of FFN551 uptake with VMAT2 inhibitor (black) or without (green) and linear fits (dashed lines). (**G**) FFN511 fluorescence intensity inside cells decreases after stimulation with serotonin (5-HT, 100 μM). Note that the decrease in the control trace is due to bleaching (n = 3, mean ± SEM (shaded)).

To visualize CA trafficking we first employed FFNs.(*28*) Neutrophils were incubated with FFN511, which has a high affinity for VMAT2 and DAT.(*21*) FFN511 localized in granular structures inside neutrophils (Fig. 1D), which is morphologically comparable to the vesicles in the adrenergic PC-12 cell line (positive control in Fig. 1D). To further verify the function of VMAT2 the uptake of FFN511 into neutrophils was observed both under normal conditions and in presence of the VMAT2 inhibitor tetrabenazine (Fig. 1E).(*33*) Unspecific uptake of FFNs has been reported and was also found in cell lines without DAT/VMAT2 (HEK and HeLa cells, Fig. S3).(*21*) Nevertheless, uptake kinetics of neutrophils under VMAT2 inhibition was slower compared to the controls (Fig. 1F), which underscores that CA uptake and packaging in neutrophils is similar to neurons and takes place within tens of seconds.

It has been long known that neutrophils can be stimulated with N-formylmethionyl-leucyl-phenylalanine (fMLP) to secret vesicular components but fMLP is not a compound that is present *in vivo*.(*34*) Platelet-derived 5-HT has been reported to modulate several neutrophil activities such as recruitment and degranulation.(*35*) Therefore, we next tested the hypothesis that 5-HT triggers exocytosis of FFN-positive vesicles. Indeed, 5-HT (100 μM) induced degranulation and release of FFN511 from neutrophils (Fig. 1G). FFN511 imaging also allowed tracking of single vesicles during 5-HT-induced exocytosis, evidenced by a stepwise decrease of fluorescence intensity (Fig. S4A).

We also employed FFN102, which is a pH-sensitive molecule with high affinity for DAT and VMAT2.(*36*) The fluorescence of FFN102 is reduced inside vesicles due to their acidic pH. Fluorescence in the FFN102 channel around the neutrophils increased after stimulation with 5-HT (Fig. S4B, C) compared to buffer, which further supports that exocytosis of vesicle content occurs.

### Serotonin triggers exocytosis of catecholamines from neutrophils

Transient release of small molecules from individual cells is difficult to capture with standard methods. Therefore, we employed a fluorescent nanosensor-based approach to directly image CA release from neutrophils with high spatiotemporal resolution as previously used for neurons.(*24, 27, 28*) For this purpose, we functionalized (6,5)-chirality enriched nIR fluorescent SWCNTs with (GT)_10_ oligonucleotides (Fig. 2A and Fig. S5A). This DNA sequence renders SWCNTs sensitive to CA. These nanosensors increased their fluorescence (in solution) by > 50 % in response to DA (100 μM) as expected (Fig. S5B).(*24, 27, 28*) Then they were homogenously physiosorbed on standard (glass) cell culture dishes (Fig. S5C). Neutrophils were then seeded on top of the nanosensors (Fig. 2B). DA (100 μM) increased the fluorescence of this thin sensor layer by ∼ 30 % whereas buffer, ionomycin (Ca^2+^ ionophore) and 5-HT (Fig. S5D) did not affect the fluorescence intensity of the immobilized nanosensors. H_2_O_2_ did also not change the fluorescence (Fig. S5E), which is important because reactive oxygen species such as H_2_O_2_ can be formed by neutrophils upon activation. These control experiments show that the nanosensor layer can specifically report local changes in CA concentration.

**Fig. 2.**
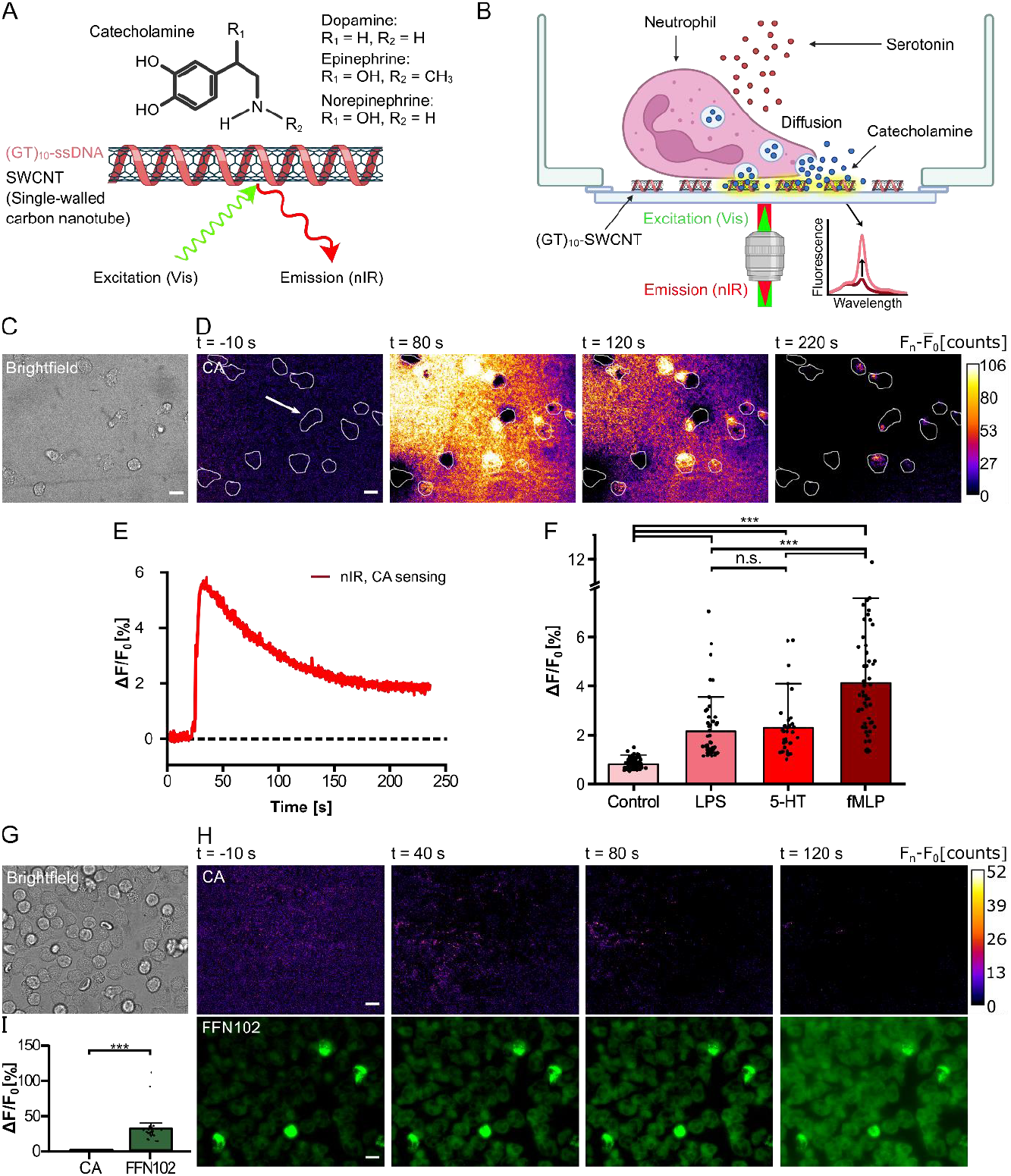
Exocytosis of catecholamines from neutrophils. (**A**) Schematic of the single-walled carbon nanotube (SWCNT) based nanosensors used to image catecholamine (CA) release. Their nIR fluorescence (980 nm) increases by CA. (**B**) Schematic of the experimental setup including immobilized nanosensors below adhering neutrophils. (**C**) Brightfield image and (**D**) nIR image sequence of neutrophils adhered on a nanosensor surface. The starting intensity is the mean pixel intensity value 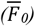 of the 10 s before addition of serotonin (5-HT). 5-HT (100 μM, t = 0) induces an increase in the fluorescence of nanosensors in localized regions corresponding to cell regions as well as everywhere by diffusion indicating exocytosis of CAs. Scale bar is 10 μm. (**E**) Fluorescence response of nanosensors to CA exocytosis from a single neutrophil after stimulation with 5-HT (around t = 20 s). The data was corrected for drift. (**F**) The fluorescence response of nanosensors (area under the cells) to CA exocytosis after stimulation with 5-HT, N-formylmethionine-leucyl-phenylalanine (fMLP), lipopolysaccharide (LPS), and a control. (n = 4, mean ± SEM, Kruskal-Wallis test, Mann-Whitney tests: p-value *** < 0.0001, each dot is a single neutrophil). (**G**) Brightfield images (cells) and (**H**) nIR image sequence (CA nanosensors, top) and green channel (FFN102, bottom) of neutrophils incubated with FFN102 before and after addition of 5-HT (100 μM). The background 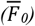 is measured 10 s before addition of 5-HT. There is no fluorescence change in the nIR, indicating no CA release but the green channel (fluorescence increase in the extracellular space) indicates FFN102 release. Scale bar is 10 μm. (**I**) Maximum fluorescence response of the nanosensors (CA release) and green channel (FFN102 release) of neutrophils stimulated with 5-HT (100 μM) after incubation with FFN102 (n = 2, mean ± SEM, unpaired t-test: p-value * < 0.05, each dot is a single neutrophil).

5-HT stimulation of neutrophils increased the nanosensor fluorescence, which indicates CA exocytosis. Hot spots of CA release are associated with cell bodies but the fast diffusion of CA quickly blurred the images (Fig. 2C, D). Release is followed by diffusion into the free extracellular space and therefore single cell (fluorescence intensity under the cell body) traces are characterized by a steep increase followed by a slow decline (Fig. 2E). We observed variation in CA release amplitudes amongst donors and experiments (Fig. S6), which shows that there is a certain heterogeneity. Other degranulation stimuli (fMLP, LPS) also induced CA exocytosis (Fig. 2F). fMLP induced release of CA could also be validated by an ELISA, when cells were pre-treated with latrunculin A (inhibits actin polymerization), which is known to increase fMLP induced exocytosis (Fig. S7).(*37*) However, only 5-HT is relevant *in vivo* and therefore we focused on this stimulus.

Neutrophils release various molecules upon activation. To exclude activation of the nanosensors by non-CA molecules, we then depleted CA in neutrophils by incubating them with FFN102 molecules. At high concentrations, FFN102s deplete VMAT2-positive vesicles from CAs and inhibit reuptake, without affecting other vesicular content.(*36*) As discussed above FFN102 fluorescence increases once released into extracellular space. FFN102-treated neutrophils were then stimulated with 5-HT and imaged simultaneously in the Vis (FFN102) and in the nIR (nanosensors) to detect endogenous CA (Fig. 2G, H). The nIR fluorescence did not increase while FFN102 fluorescence significantly increased, indicating exocytosis of FFN102 and depletion of CA (Fig. 2I). These results underline that the nanosensors specifically report fast and transient CA release and VMAT2 positive vesicles play the central role.

### Serotonin triggers intracellular Ca^2+^ signals and catecholamine exocytosis

Calcium signaling plays an important role in key neutrophil functions.(*38*) For example, exocytosis of gelatinase, specific, and azurophilic granules in neutrophils is triggered by transient elevation in the intracellular calcium concentration. To better understand the processes accompanied by the 5-HT-induced CA exocytosis we used the Ca^2+^ indicator Fluo-4 AM.(*39*) Ionomycin-induced Ca^2+^ influx was employed as a positive control (Fig. S8). Indeed, 5-HT but not DA induced a significant spike in the intracellular concentration of Ca^2+^ (>50 % increase) (Fig. 3A, B). We employed the above-mentioned nanosensor-based sensing strategy to visualize exocytosis of CAs after stimulation with 5-HT together with Ca^2+^ imaging (Fig. 3C, D). Neutrophils were seeded again on a nanosensor surface, and the Fluo-4 AM fluorescence was imaged in the visible channel (GFP), while CA exocytosis was recorded in the nIR channel (5 frames/s). 5-HT increased the intracellular Ca^2+^ concentration, followed by a delayed (around 10-20 s) CA release (Fig. 3C, D). This time scale is different from neuronal DA release, which typically happens within ms, suggesting that there are no primed vesicles in neutrophils as in neurons.(*28*) A numerical diffusion simulation of the DA concentration, as an exemplary CA, in the immediate vicinity of a DA-releasing cell was performed to better understand the obtained traces (Fig. 3D). The simulated cell released DA over the course of 80 s from several vesicles (Fig. 3E and Tables S4, S5). Importantly, the measurable fluorescence signals are not the concentration but rather the convolution between concentration and sensor kinetics. Therefore, we also simulated the corresponding amount of sensor-bound DA (Fig. 3F) accounting for on-off kinetics (Table S4, Table S5). The qualitative similarity to Fig. 3D indicates that CAs are released from several vesicles over several 10s of seconds around the cell as opposed to other simulated scenarios such as a single release event (Fig. S9, Table S5).

**Fig. 3.**
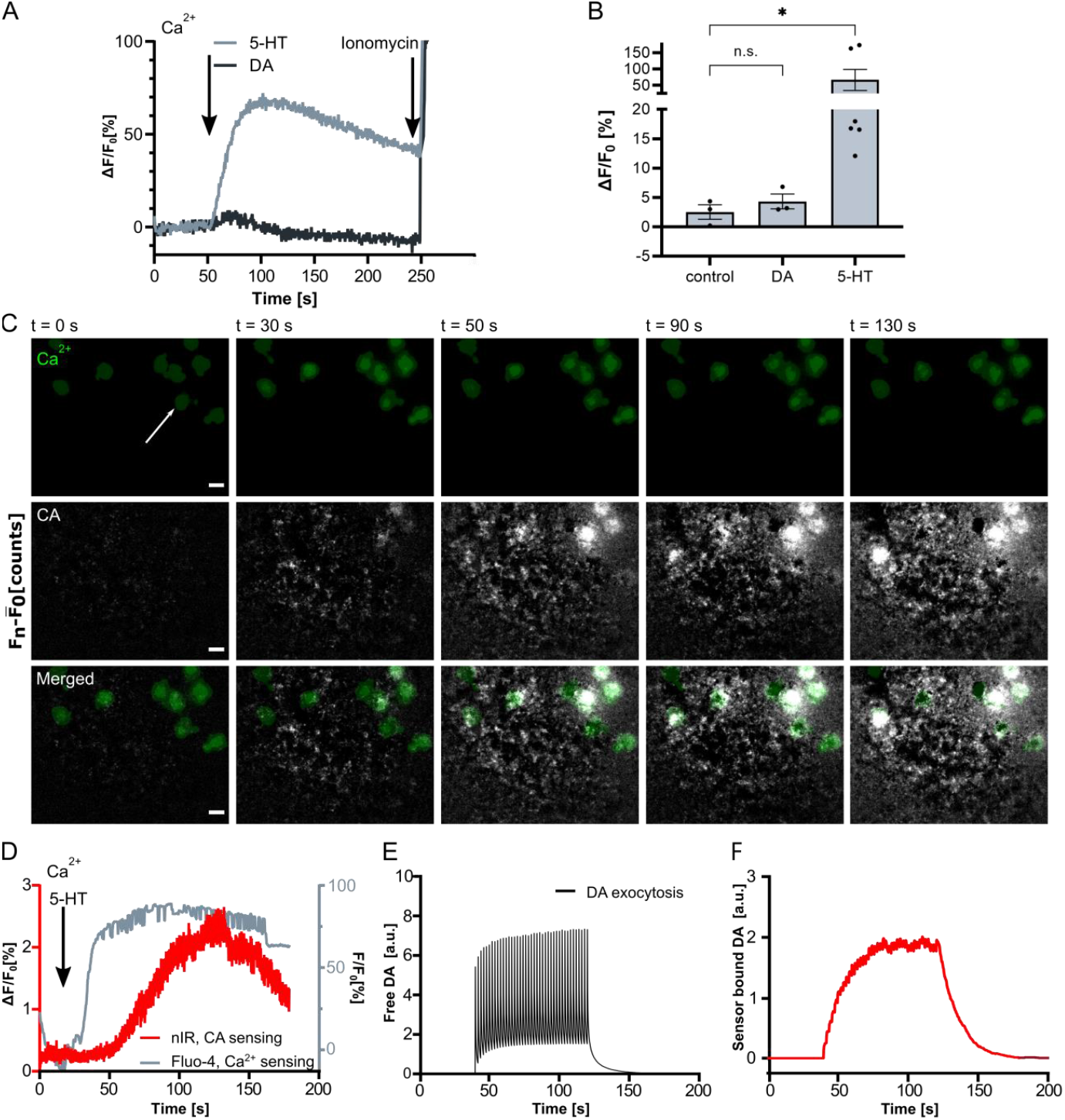
Serotonin triggers intracellular Ca^2+^ signals and catecholamine exocytosis. (**A**) Fluorescence signal of neutrophils incubated with Fluo-4 AM over time. Serotonin (5-HT) or dopamine (DA) (100 μM) is added at t = 50 s and ionomycin (5 μM) at t = 250 s. (**B**) Maximum fluorescence increase (calcium concentration change) for DA and 5-HT (n ≥ 3 donors, mean ± SEM, Kruskal-Wallis test with Dunn’s multiple comparisons test: p value ** < 0.001, each dot is a single experiment). (**C**) Neutrophils are adhered on nanosensor-coated glass slides. Fluo-4 AM (Ca^2+^) and nIR nanosensor channel (CA) before and after stimulation with 5-HT (100 μM) at t = 0. All scale bars are 10 μm. (**D**) CA (nIR nanosensors, red) and Ca^2+^ (Fluo-4, grey) time trace of an exemplary neutrophil (arrow in (D)). Fluorescence changes are the normalized intensity change under the cell outline (from the brightfield contour). (**E**) CA release and diffusion simulation: Simulated DA concentration in the area of a cell (diameter = 10 μm), after DA is released from several vesicles at the cell membrane in release events every 2 s over the course of 80 s (from t = 40 s to t = 120 s). (**F**) Corresponding simulated nanosensor fluorescence response, which is a convolution of concentration (E) and on-off sensor kinetics.

### Serotonin receptors are required and platelet-neutrophil interactions trigger CA release

To investigate whether CA exocytosis induced by 5-HT is 5-HT receptor mediated, we treated neutrophils with the 5-HT2 selective antagonist ketanserin.(*40*) After neutrophils adhered to the nanosensor-coated surface, ketanserin (100 μM) was added for 2-5 min before stimulating them. Ketanserin-treated neutrophils did not release CA when exposed to 5-HT (Fig. 4A, Fig. S10A, B). However, ionomycin still increased fluorescence at localized regions around cells corresponding to CA release comparable to that of 5-HT (Fig. S10A - C). The 5-HT receptor agonist 2-[(3-chlorophenyl)methoxy]-6-(1-piperazinyl)-pyrazine (CP809) also induced exocytosis of CA but with slower kinetics (Fig. 4B, C, Fig. S10D, E). These results indicate that at least 5-HT2 engagement is necessary for CA exocytosis in neutrophils.

**Fig. 4.**
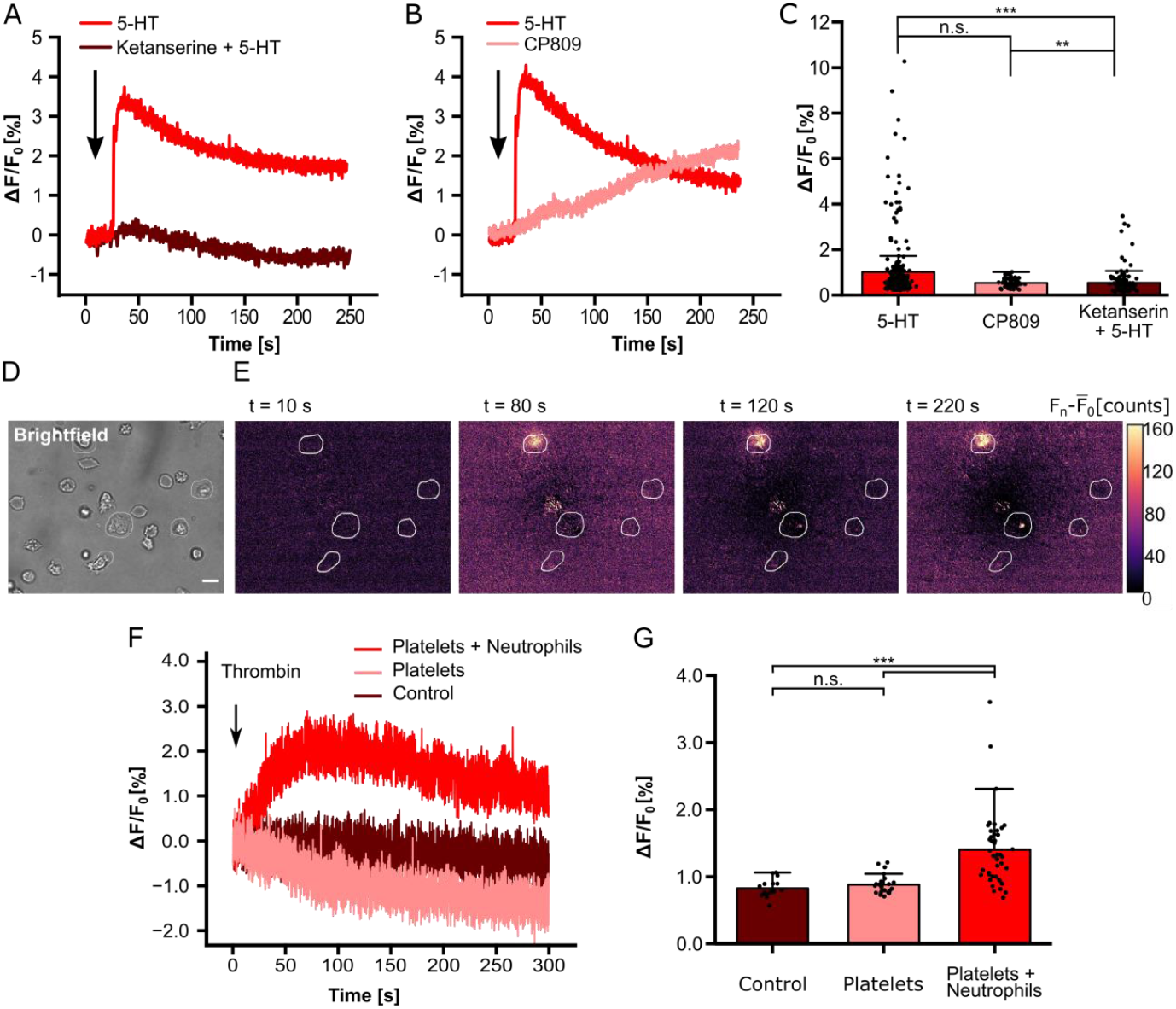
Catecholamine exocytosis is serotonin receptor dependent and triggered by activated platelets. (**A**) Nanosensor signal from exemplary neutrophil stimulated with serotonin (5-HT, 100 μM) and treated with and without 5-HT antagonist ketanserin (100 μM). (**B**) Nanosensor signal from exemplary neutrophil stimulated with serotonin (5-HT) or 5-HT agonist CP809. (**C**) Maximum nanosensor fluorescence increase of neutrophils after stimulation with 5-HT (100 μM), treated with ketanserin (100 μM) or CP809 (1 μM). (n ≥ 5, mean ± SEM, Kruskal-Wallis test and Mann-Whitney tests: p-value *** < 0.001, ** < 0.01, each dot is a single neutrophil). (**D**) Brightfield image and (**E**) nIR image sequence (right) of neutrophils and platelets adhered on a nanosensor-coated glass surface. nIR images are recorded before and after stimulation of platelets with thrombin (1 Unit/mL). The outlines indicate the shapes of neutrophils. (**F**) Exemplary fluorescence responses of nanosensors to neutrophil + thrombin + adhered platelets, thrombin + adhered platelets and thrombin (control). (**G**) Maximum fluorescence increase of nanosensors for a control without cells, adhered platelets + thrombin, and neutrophil + adhered platelets + thrombin (n = 3, mean ± SEM, Kruskal-Wallis test and Mann-Whitney tests: p-value * < 0.5, *** <0.001, each data point corresponds to a single neutrophil).

Platelets store most of the 5-HT in humans and therefore we studied platelet-neutrophil interactions by seeding platelets and neutrophils together on nanosensor-coated surfaces. Neutrophils exposed to platelets + thrombin released CA (Fig. 4D - G). In contrast, thrombin alone as well as thrombin + platelets did not change the fluorescence (Fig. 4F). This shows that the interaction of activated platelets and neutrophilic granulocytes activates exocytosis of CA by neutrophils. To exclude the impact of thrombin on neutrophils neutrophils were inhibited with ketanserin and showed no effect upon platelet trigger with thrombin (Fig. S11).

### Paracrine modulatory role of catecholamines on formation of neutrophil extracellular traps and platelet aggregation

CAs have an affinity to bind a large variety of receptors including different DA and adrenergic receptors. The expression of beta-adrenergic receptors on neutrophils has long been established (*17*) and it is known that these receptors bind not only E and NE but DA as well.(*41*) Therefore, the expression of the adrenergic receptors ADRB1 and ADRB2 as exemplary CA receptors was verified using immunofluorescence staining (Fig. 5A) on the protein level and qPCR on RNA level (Fig. 5B). Interestingly, ADRB1 expression is increased in neutrophils incubated with 50 ng/mL LPS for two hours (Fig. S12), pointing to a role of ADRB1 in stimulated cells. It also suggests that the cells could change their responsiveness to CAs by changing the expression pattern of CA receptors. Therefore, we examined the effect of various monoamines (E, NE, DA, 5-HT) on the rate of NETosis with and without stimulation with phorbol-12-myristate-13-acetate (5 nM PMA), a commonly used stimulus of NET formation. All monoamines reduced NET formation compared to the control group in a concentration-dependent manner (Fig. 5C, Fig. S13). Therefore, the CAs seem to have a direct effect on the NETosis rate of neutrophilic granulocytes. In the case of 5-HT, this may be caused directly or indirectly by stimulating CA exocytosis.

**Fig. 5.**
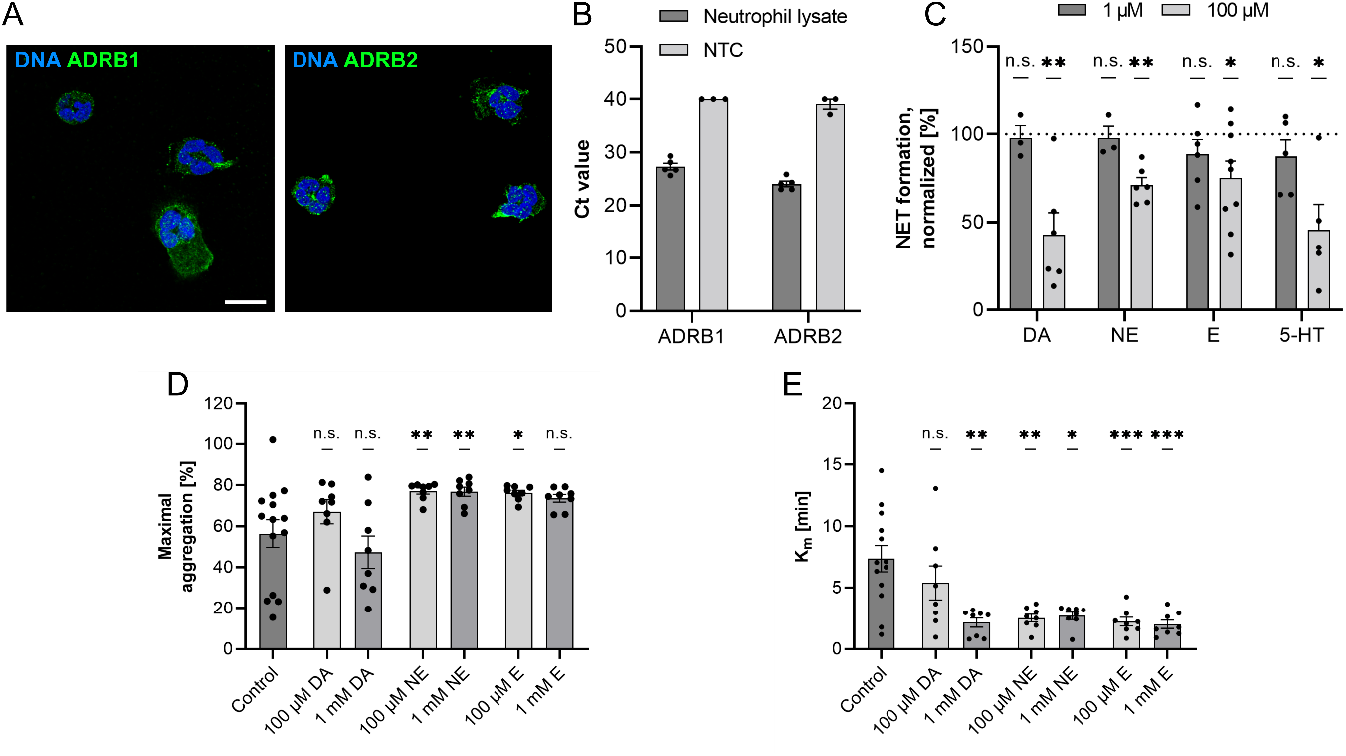
Paracrine catecholamine signaling impacts NETosis and thrombosis. (**A**) Neutrophils stained for β1-adrenergic receptor (ADRB1, green) or β2-adrenergic receptor (ADRB2, green) and DNA (blue). Scale bar is 10 μm. (**B**) Both receptors are found at the transcriptional level as determined by qPCR (n = 5 and n = 3, mean ± SEM, NTC: no template control). (**C**) Effect of CAs as well as serotonin on NETosis in neutrophils stimulated with 5 nM PMA. PMA stimulated controls = 100 %, (n ≥ 3, mean ± SEM, one-sample t-test, or one-sample Wilcoxon test: p-value * < 0.05, ** < 0.01). (**D, E**) Platelets are stimulated with thrombin in combination with CAs or thrombin alone (control) to determine the aggregation maximum (D) or the aggregation speed (E) using aggregometry. K_m_ represents the time until half-maximal aggregation). n ≥ 8, mean ± SEM, Kruskal-Wallis test with uncorrected Dunn’s test: p-value * < 0.05, ** < 0.01).

Next, we investigated whether CAs released by neutrophils affect platelet function. To this end, we quantified thrombin-induced platelet aggregation using the well-established method of optical aggregometry.(*42*) Platelet aggregation is an important part of thrombus formation in response to blood vessel injury as well as in thrombosis. NE and E but not DA increased the overall maximal aggregation response of thrombin-stimulated platelets (Fig. 5D) and all CAs increased the aggregation speed as evidenced by a smaller K_m_ value (time until half-maximal aggregation). NE and E elicited this platelet response at lower concentrations than DA (Fig. 5E).

## Discussion

There is growing evidence that catecholamines (CAs) such as dopamine (DA), epinephrine (E), or norepinephrine (NE) affect the function of immune cells, which can have severe consequences to patients receiving therapeutic doses of them during sepsis or other life-threatening events (*43*–*45*). However, if and how immune cells synthesize, store, release, and respond to transient, localized CA signals remained elusive.

We aimed to study the catecholaminergic system and its functional role in the most abundant type of immune cells (neutrophils). First, we wanted to find out whether the dopaminergic machinery known from neurons is present. We confirmed the finding (*46*) that TH, the rate-limiting enzyme for CA synthesis, is expressed in neutrophils. Additionally, we found the CA transporters DAT in the membrane and VMAT2 in azurophilic granules as well as CA degradation enzymes (Fig. 1A–C). DA and to a lesser extent E and NE were measured inside neutrophils (Fig. 1D), as well as metabolic (degradation) products of CAs. Thus, neutrophils can take up, store and process CAs.

Additionally, we studied CA uptake and release by using FFNs (Fig. 1E) that are a well-recognized tool for studying neuronal cells but have to our knowledge never been employed in neutrophils or other immune cells. (*21, 36*) Our results reveal that VMAT2 is involved in CA uptake (Fig. 1F, G). The very fast FFN uptake by neutrophils within seconds suggest that neutrophils could play an important role in systemic and/or local clearance of CAs.

Interestingly, we were able to identify serotonin (5-HT) as a trigger for CA exocytosis (Fig. 1G). 5-HT is stored and released by blood platelets and plays a role in degranulation of neutrophilic granulocytes.(*35*) This suggests a possible functional feedback loop between platelets and neutrophils. Taking into account that FFNs only indirectly prove CA exocytosis, we next made use of the much higher spatiotemporal resolution of nIR fluorescent nanosensors, specifically tailored for CA detection (*47*), and observed fast and transient CA exocytosis events (Fig. 2A–F).

CA concentrations in vesicles in neurons are thought to range up to 300 mM.(*48*) However, CA exocytosis causes a local peak of CA concentration that rapidly decays due to diffusion within a few hundred milliseconds. Thus, only a method with a high spatiotemporal resolution is able to resolve CA release (Fig. 2) and other methods will most likely fail to capture these events. Our observation of CA exocytosis from neutrophils is further supported by experiments with another class of FFN (FFN102), which increase their fluorescence when released into extracellular space following serotonin stimulation (Fig. 2G–I).

Several studies indicate that Ca^2+^ mobilization plays an important role in all types of neutrophil degranulation.(*49*) To further elucidate the mechanism of CA release, we therefore imaged both Ca^2+^ and CA in a custom-built dual nIR/Vis fluorescence microscope. We found that 5-HT induces an intracellular Ca^2+^ signal followed by CA release (Fig. 3A–D). Interestingly, this process takes much longer (typically > 10 s) than CA release in neurons (milliseconds) suggesting that the vesicles need more time to successfully fuse with the membrane. Indeed, CA release from neutrophils can be modeled *in silico* and supports the notion of relatively slow release over the time course of several tens of seconds (typically 10 s – 60 s) (Fig. 3E, F). We also found that 5-HT stimulation requires at least 5-HT2 receptors (Fig. 4A, C).

To further investigate the putative interplay between neutrophils and platelets in a more physiological scenario, we brought neutrophils in contact with platelets and activated them with thrombin, which is known to induce 5-HT release.(*50*) Indeed, this interaction triggered CA release (Fig. 4D–G), which establishes an immunomodulatory link (via serotonin ad CA) between platelets and neutrophils. This link appears especially relevant as we also provide evidence that neutrophil behavior, particular NET formation can be modulated by CA (Fig. 5C), corroborating previous reports of an inhibitory effect of CA on proinflammatory macrophage and dendritic cell functions.(*15*) Previous studies found involvement of both alpha- and beta-adrenergic receptors in the catecholaminergic signaling in the peripheral immune system.(*16, 51*) Indeed, we also found adrenergic receptors ADRB1 and 2 to be prominently expressed in neutrophils (Fig. 5A, B, S11).

Receptors often exhibit a cross-reactivity with several, similar molecules. For example, the β1- and β2-adrenergic receptors (ADRB1 and ADRB2, Fig. 5A, B) respond to E and NE with K_d_ values in the low micromolar range but also to DA with K_d_ values in the higher micromolar range.(*41*) Additionally, kinetics and local concentrations play an equally important role. Therefore, given the expression of DA receptors (*14*) on neutrophils there seem to be multiple receptors through which CAs can impact neutrophils. An alternative mechanism could be linked to the redox properties of CA.

Moreover, CAs not only impact neutrophil behavior but CA sensing has direct effects on platelet clotting, increasing maximum platelet aggregation as well as clotting speed, indicating a role for neurotransmitter-driven platelet-neutrophil interactions in thromboinflammation, a process in which neutrophils, platelets and NETs are known to play a role. One may speculate that during thromboinflammation, activated neutrophils propagate platelet aggregation via CAs and other factors, which in turn leads to serotonin release, providing a feed-forward loop for CA secretion (Fig. 6). Thus, locally released CAs affects surrounding cells, possibly in an autocrine/paracrine way.

**Fig. 6.**
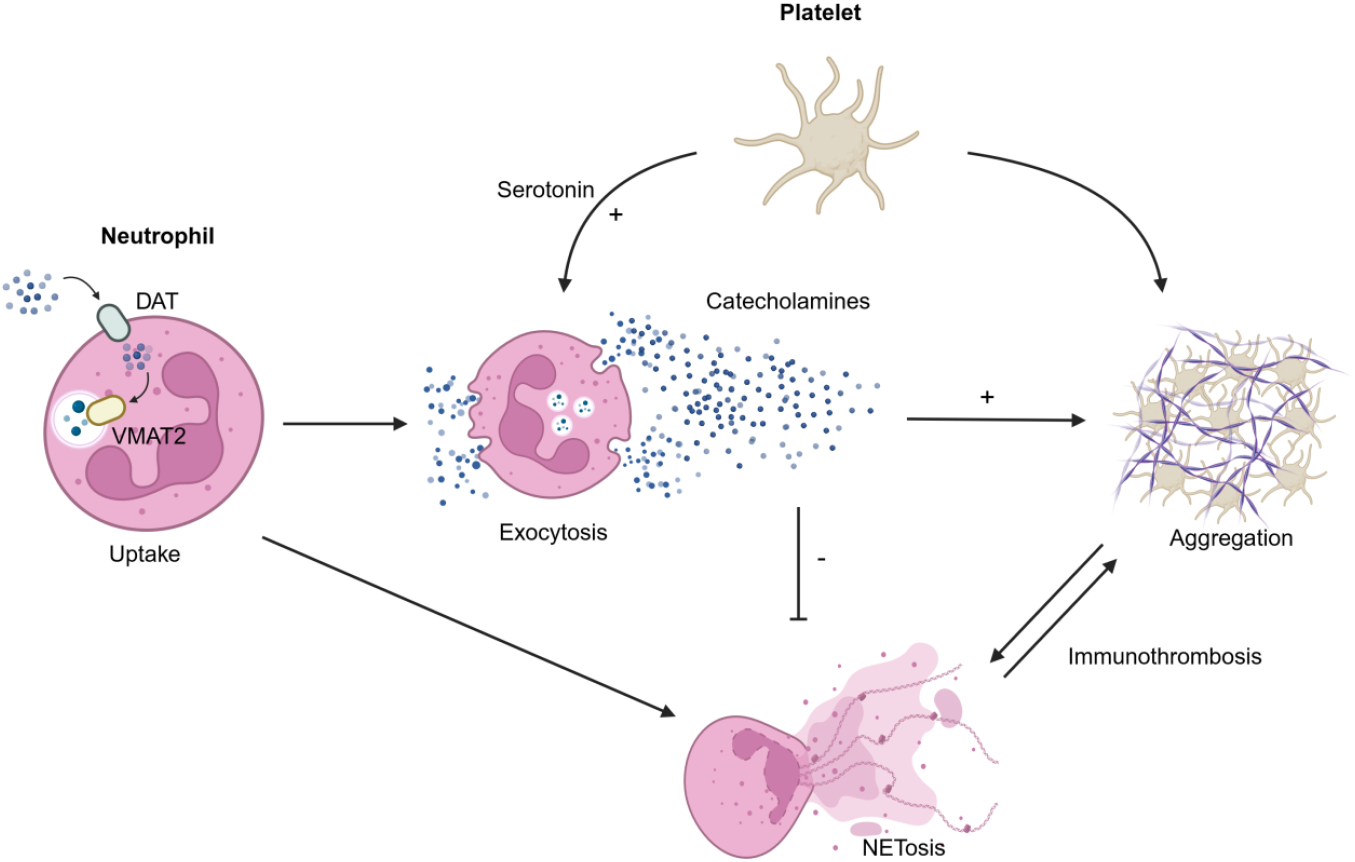
Model for catecholamine-mediated immunomodulatory feedback loop between neutrophils and platelets. Neutrophils synthesize or take up catecholamines (CAs) via DAT and package them into vesicles using VMAT2. Serotonin released by activated platelets triggers exocytosis of CAs, which enhances platelet aggregation and inhibits NETosis.

Of note, NE infusions, which are frequently used in intensive care medicine for septic shock, impair the immune defense through anti-inflammatory effects and a change in the systemic CA level, both regulating and restricting the immune response.(*52*)

All in all, our work shows that CA signaling creates a (negative) feedback loop to modulate neutrophil functions (Fig. 6). Given its local/paracrine nature, it depends on the proximity between the neutrophils as well as platelets. In a broader context, this is very similar to volume transmission of CA in the brain.(*48*) The neutrophils have multiple functions regarding Cas (synthesis, uptake, degradation, release, detection) and thus can contribute to a complicated regulation system. Importantly, the biological implications of local, rapid and transient CA release from neutrophils are likely to differ from what is known from massive systemic application of i.e. NE during sepsis, as this autocrine/paracrine loop could be used to adjust immune functions and platelet aggregation in a much more refined manner.

Subpopulations of neutrophils could also respond in different ways, which corresponds to plasticity in the neuronal system. Despite originally perceiving neutrophils as a terminally differentiated and homogenous population, several subpopulations of neutrophils with distinct molecular biomarkers have been discovered.(*53*) Different subtypes of neutrophils may have different CA content and different levels of CA release as seen in the variation between different donors and cells (Fig. S6).

Our findings also raise the question if there are more CA functions e.g. between immune cells and endothelial cells and if they are of similar importance to the immune system as to the neuronal system. A major difference is that immunological interactions between cells are typically transient and not hard-wired as in the brain. Therefore, CA functions likely also depend on cellular proximity and adhesion status.

## Conclusion

Catecholamines (CAs) are well-known for their role as neurotransmitters in the neuronal system and as hormones. We provide evidence that neutrophils have the machinery for CA trafficking and show their impact on different functions of neutrophils and platelets. The fast CA release and uptake events could be captured by a CA-sensitive nanosensor imaging approach. Overall, we propose that the immune system uses CAs as immunomodulators through rapid spatiotemporal concentration changes.

## Acknowledgments

We sincerely thank everyone for their contribution in consistently performing blood sample collections essential to this study.

## Funding

Deutsche Forschungsgemeinschaft (DFG, German Research Foundation) under Germany’s Excellence Strategy – EXC 2033 – 390677874 – RESOLV (S.K.)”Center for Solvation Science ZEMOS” funded by the German Federal Ministry of Education and Research BMBF and by the Ministry of Culture and Research of Nord Rhine-Westphalia (S.K.). This work was funded by the DFG (L.E., S.K.).

Author contributions

Conceptualization: SK, LE

Methodology: JM, AS, MD, SJ, IB, SK, LE

Investigation: JM, AS, MD, BH, MS, SJ, SS, EN, SKa, LK

Visualization: JM, AS, MD

Funding acquisition: SK, LE

Project administration: SK, LE

Supervision: SK, LE

Writing – original draft: MD, MS, AS, JM, SK, LE

Writing – review & editing: JM, AS, SK, LE, EN, IB

## Diversity, equity, ethics, and inclusion [optional]

This study was approved by the Ethics Committee Westfalen-Lippe (approval number 2021-657-f-S). Before donating blood, fully informed consent of each donor was obtained.

## Data and materials availability

All data are available in the main text or the supplementary materials. Raw data will be uploaded to a repository once the paper has been accepted.

## References

1. K. H. Lee, A. Kronbichler, D. D.-Y. Park, Y. Park, H. Moon, H. Kim, J. H. Choi, Y. Choi, S. Shim, I. S. Lyu, Neutrophil extracellular traps (NETs) in autoimmune diseases: a comprehensive review. Autoimmun Rev 16, 1160–1173 (2017).

2. M. Siwicki, P. Kubes, Neutrophils in host defense, healing, and hypersensitivity: Dynamic cells within a dynamic host. J Allergy Clin Immunol 151, 634–655 (2023).

3. J. S. Gauer, R. A. Ajjan, R. A. S. Ariëns, Platelet-Neutrophil Interaction and Thromboinflammation in Diabetes: Considerations for Novel Therapeutic Approaches. J Am Heart Assoc 11 (2022).

4. A. Caudrillier, K. Kessenbrock, B. M. Gilliss, J. X. Nguyen, M. B. Marques, M. Monestier, P. Toy, Z. Werb, M.R. Looney, Platelets induce neutrophil extracellular traps in transfusion-related acute lung injury. J Clin Invest 122, 2661–2671 (2012).

5. C. Liu, P. Goel, P. S. Kaeser, Spatial and temporal scales of dopamine transmission. Nature Reviews Neuroscience 2021 22:6 22, 345–358 (2021).

6. M. Kreft, J. Jorgačevski, N. Vardjan, R. Zorec, Unproductive exocytosis. J Neurochem 137, 880–889 (2016).

7. T. C. Südhof, Neurotransmitter release: the last millisecond in the life of a synaptic vesicle. Neuron 80, 675–690 (2013).

8. T. C. Südhof, The synaptic vesicle cycle. Annu Rev Neurosci 27, 509–547 (2004).

9. M. R. Logan, S. O. Odemuyiwa, R. Moqbel, Understanding exocytosis in immune and inflammatory cells: The molecular basis of mediator secretion. Journal of Allergy and Clinical Immunology 111, 923–932 (2003).

10. I. Papa, D. Saliba, M. Ponzoni, S. Bustamante, P. F. Canete, P. Gonzalez-Figueroa, H. A. McNamara, S. Valvo, M. Grimbaldeston, R. A. Sweet, H. Vohra, I. A. Cockburn, M. Meyer-Hermann, M. L. Dustin, C. Doglioni, C.G. Vinuesa, TFH-derived dopamine accelerates productive synapses in germinal centres. Nature 547, 318–323 (2017).

11. N. R. Musso, S. Brenci, M. Setti, F. Indiveri, G. Lotti, Catecholamine content and in vitro catecholamine synthesis in peripheral human lymphocytes. J Clin Endocrinol Metab 81, 3553–3557 (1996).

12. P. J. Gaskill, L. Carvallo, E. A. Eugenin, J. W. Berman, Characterization and function of the human macrophage dopaminergic system: implications for CNS disease and drug abuse. J Neuroinflammation 9, 203 (2012).

13. E. Rönnberg, G. Calounova, G. Pejler, Mast cells express tyrosine hydroxylase and store dopamine in a serglycin-dependent manner. Biol Chem 393, 107–112 (2012).

14. F. McKenna, P. J. McLaughlin, B. J. Lewis, G. C. Sibbring, J. A. Cummerson, D. Bowen-Jones, R. J. Moots, Dopamine receptor expression on human T- and B-lymphocytes, monocytes, neutrophils, eosinophils and NK cells: A flow cytometric study. J Neuroimmunol 132, 34–40 (2002).

15. A. Scanzano, M. Cosentino, Adrenergic regulation of innate immunity: a review. Front Pharmacol 6 (2015).

16. D. Sharma, J. D. Farrar, Adrenergic regulation of immune cell function and inflammation. Semin Immunopathol 42, 709–717 (2020).

17. S. P. Galant, S. Allred, Binding and functional characteristics of beta adrenergic receptors in the intact neutrophil. J Lab Clin Med 98, 227–237 (1981).

18. F. Marino, A. Scanzano, L. Pulze, M. Pinoli, E. Rasini, A. Luini, R. Bombelli, M. Legnaro, M. de Eguileor, M. Cosentino, β(2) -Adrenoceptors inhibit neutrophil extracellular traps in human polymorphonuclear leukocytes. J Leukoc Biol 104, 603–614 (2018).

19. A. Scanzano, L. Schembri, E. Rasini, A. Luini, J. Dallatorre, M. Legnaro, R. Bombelli, T. Congiu, M. Cosentino, F. Marino, Adrenergic modulation of migration, CD11b and CD18 expression, ROS and interleukin-8 production by human polymorphonuclear leukocytes. Inflamm Res 64, 127–135 (2015).

20. F. Sun, J. Zeng, M. Jing, J. Zhou, J. Feng, S. F. Owen, Y. Luo, F. Li, H. Wang, T. Yamaguchi, Z. Yong, Y. Gao, W. Peng, L. Wang, S. Zhang, J. Du, D. Lin, M. Xu, A. C. Kreitzer, G. Cui, Y. Li, A Genetically Encoded Fluorescent Sensor Enables Rapid and Specific Detection of Dopamine in Flies, Fish, and Mice. Cell 174, 481-496.e19 (2018).

21. N. G. Gubernator, H. Zhang, R. G. W. Staal, E. V. Mosharov, D. B. Pereira, M. Yue, V. Balsanek, P. A. Vadola, B. Mukherjee, R. H. Edwards, D. Sulzer, D. Sames, Fluorescent false neurotransmitters visualize dopamine release from individual presynaptic terminals. Science 324, 1441–1444 (2009).

22. J. Ackermann, J. T. Metternich, S. Herbertz, S. Kruss, Biosensing with fluorescent carbon nanotubes. Angewandte Chemie International Edition 61, e202112372 (2022).

23. M. J. O’Connell, S. H. Bachilo, C. B. Huffman, V. C. Moore, M. S. Strano, E. H. Haroz, K. L. Rialon, P. J. Boul, W. H. Noon, C. Kittrell, J. Ma, R. H. Hauge, R. B. Weisman, R. E. Smalley, Band gap fluorescence from individual single-walled carbon nanotubes. Science (1979) 297, 593–596 (2002).

24. S. Kruss, M. P. Landry, E. Vander Ende, B. M. A. Lima, N. F. Reuel, J. Zhang, J. Nelson, B. Mu, A. Hilmer, M. Strano, Neurotransmitter detection using corona phase molecular recognition on fluorescent single-walled carbon nanotube sensors. J Am Chem Soc 136, 713–724 (2014).

25. H. Wu, R. Nißler, V. Morris, N. Herrmann, P. Hu, S. J. Jeon, S. Kruss, J. P. Giraldo, Monitoring Plant Health with Near-Infrared Fluorescent H2O2 Nanosensors. Nano Lett 20, 2432–2442 (2020).

26. G. Bisker, J. Dong, H. D. Park, N. M. Iverson, J. Ahn, J. T. Nelson, M. P. Landry, S. Kruss, M. S. Strano, Protein-targeted corona phase molecular recognition. Nat Commun 7, 1–14 (2016).

27. S. Kruss, D. P. Salem, L. Vuković, B. Lima, E. Vander Ende, E. S. Boyden, M. S. Strano, High-resolution imaging of cellular dopamine efflux using a fluorescent nanosensor array. Proc Natl Acad Sci U S A 114, 1789–1794 (2017).

28. S. Elizarova, A. A. Chouaib, A. Shaib, B. Hill, F. Mann, N. Brose, S. Kruss, J. A. Daniel, A fluorescent nanosensor paint detects dopamine release at axonal varicosities with high spatiotemporal resolution. Proc Natl Acad Sci U S A 119, e2202842119 (2022).

29. M. Dinarvand, E. Neubert, D. Meyer, G. Selvaggio, F. A. Mann, L. Erpenbeck, S. Kruss, Near-Infrared Imaging of Serotonin Release from Cells with Fluorescent Nanosensors. Nano Lett 19 (2019).

30. E. N. Pothos, K. E. Larsen, D. E. Krantz, Y. Liu, J. W. Haycock, W. Setlik, M. D. Gershon, R. H. Edwards, D. Sulzer, Synaptic vesicle transporter expression regulates vesicle phenotype and quantal size. Journal of Neuroscience 20, 7297–7306 (2000).

31. M. J. Nirenberg, R. A. Vaughan, G. R. Uhl, M. J. Kuhar, V. M. Pickel, The dopamine transporter is localized to dendritic and axonal plasma membranes of nigrostriatal dopaminergic neurons. The Journal of neuroscience 16, 436 (1996).

32. D. M. Omiatek, A. J. Bressler, A. S. Cans, A. M. Andrews, M. L. Heien, A. G. Ewing, The real catecholamine content of secretory vesicles in the CNS revealed by electrochemical cytometry. Scientific Reports 2013 3:1 3, 1–6 (2013).

33. S. I. Støve, Å. A. Skjevik, K. Teigen, A. Martinez, Inhibition of VMAT2 by β2-adrenergic agonists, antagonists, and the atypical antipsychotic ziprasidone. Communications Biology 2022 5:1 5, 1–14 (2022).

34. N. R. Jog, M. J. Rane, G. Lominadze, G. C. Luerman, R. A. Ward, K. R. McLeish, The actin cytoskeleton regulates exocytosis of all neutrophil granule subsets. Am J Physiol Cell Physiol 292 (2007).

35. M. Mauler, N. Herr, C. Schoenichen, T. Witsch, T. Marchini, C. Härdtner, C. Koentges, K. Kienle, V. Ollivier, M. Schell, L. Dorner, C. Wippel, D. Stallmann, C. Normann, H. Bugger, P. Walther, D. Wolf, I. Ahrens, T. Lämmermann, B. Ho-Tin-Noe, K. Ley, C. Bode, I. Hilgendorf, D. Duerschmied, Platelet Serotonin Aggravates Myocardial Ischemia/Reperfusion Injury via Neutrophil Degranulation. Circulation 139, 918–931 (2019).

36. P. C. Rodriguez, D. B. Pereira, A. Borgkvist, M. Y. Wong, C. Barnard, M. S. Sonders, H. Zhang, D. Sames, D. Sulzer, Fluorescent dopamine tracer resolves individual dopaminergic synapses and their activity in the brain. Proceedings of the National Academy of Sciences 110, 870–875 (2013).

37. N. R. Jog, M. J. Rane, G. Lominadze, G. C. Luerman, R. A. Ward, K. R. McLeish, The actin cytoskeleton regulates exocytosis of all neutrophil granule subsets. Am J Physiol Cell Physiol 292 (2007).

38. J. Hann, J.-L. Bueb, F. Tolle, S. Bréchard, Calcium signaling and regulation of neutrophil functions: still a long way to go. J Leukoc Biol 107, 285–297 (2020).

39. K. R. Gee, K. A. Brown, W. N. U. Chen, J. Bishop-Stewart, D. Gray, I. Johnson, Chemical and physiological characterization of fluo-4 Ca2+-indicator dyes. Cell Calcium 27, 97–106 (2000).

40. W. C. Sessa, K. M. Mullane, Release of a neutrophil-derived vasoconstrictor agent which augments platelet-induced contractions of blood vessels in vitro. Br J Pharmacol 99, 553 (1990).

41. J. G. Baker, The selectivity of β-adrenoceptor agonists at human β _1_-, β _2_- and β _3_-adrenoceptors. Br J Pharmacol 160, 1048–1061 (2010).

42. J. Le Blanc, F. Mullier, C. Vayne, M. Lordkipanidzé, Advances in Platelet Function Testing-Light Transmission Aggregometry and Beyond. J Clin Med 9, 1–17 (2020).

43. K. C. L. Torres, L. R. V Antonelli, A. L. S. Souza, M. M. Teixeira, W. O. Dutra, K. J. Gollob, Norepinephrine, dopamine and dexamethasone modulate discrete leukocyte subpopulations and cytokine profiles from human PBMC. J Neuroimmunol 166, 144–157 (2005).

44. Y. Yan, W. Jiang, L. Liu, X. Wang, C. Ding, Z. Tian, R. Zhou, Dopamine Controls Systemic Inflammation through Inhibition of NLRP3 Inflammasome. Cell 160, 62–73 (2015).

45. R. F. Stolk, E. van der Pasch, F. Naumann, J. Schouwstra, S. Bressers, A. E. van Herwaarden, J. Gerretsen, R. Schambergen, M. M. Ruth, J. G. van der Hoeven, H. van Leeuwen, P. Pickkers, M. Kox, Norepinephrine dysregulates the immune response and compromises host defense during sepsis. Am J Respir Crit Care Med 202, 830–842 (2020).

46. M. A. Flierl, D. Rittirsch, B. A. Nadeau, J. V. Sarma, D. E. Day, A. B. Lentsch, M. S. Huber-Lang, P. A. Ward, Upregulation of phagocyte-derived catecholamines augments the acute inflammatory response. PLoS One 4, e4414 (2009).

47. F. A. Mann, N. Herrmann, D. Meyer, S. Kruss, Tuning selectivity of fluorescent carbon nanotube-based neurotransmitter sensors. Sensors (Switzerland) 17, 1521 (2017).

48. S. J. Cragg, M. E. Rice, DAncing past the DAT at a DA synapse. Trends Neurosci 27, 270–277 (2004).

49. K. Káldi, J. Szeberényi, B. K. Rada, P. Kovács, M. Geiszt, A. Mócsai, E. Ligeti, Contribution of phopholipase D and a brefeldin A-sensitive ARF to chemoattractant-induced superoxide production and secretion of human neutrophils. J Leukoc Biol 71, 695–700 (2002).

50. L. F. Brass, Thrombin and Platelet Activation. Chest 124, 18S–25S (2003).

51. S. Chhatar, G. Lal, Role of adrenergic receptor signalling in neuroimmune communication. Current Research in Immunology 2, 202 (2021).

52. J. Bakker, E. Kattan, D. Annane, R. Castro, M. Cecconi, D. De Backer, A. Dubin, L. Evans, M. N. Gong, O. Hamzaoui, C. Ince, B. Levy, X. Monnet, G. A. O. Tascón, M. Ostermann, M. R. Pinsky, J. A. Russell, B. Saugel, T. W. L. Scheeren, J. L. Teboul, A. V. Baron, J. L. Vincent, F. G. Zampieri, G. Hernandez, Current practice and evolving concepts in septic shock resuscitation. Intensive Care Med 48, 148–163 (2022).

53. X. Xie, Q. Shi, P. Wu, X. Zhang, H. Kambara, J. Su, H. Yu, S.-Y. Park, R. Guo, Q. Ren, S. Zhang, Y. Xu, L. E. Silberstein, T. Cheng, F. Ma, C. Li, H. R. Luo, Single-cell transcriptome profiling reveals neutrophil heterogeneity in homeostasis and infection. Nat Immunol 21, 1119–1133 (2020).

54. S. Elizarova, Resolving Dopamine Secretion at Individual Varicosities using Carbon Nanotube-Based Optical Dopamine Nanosensors. doi: 10.53846/GOEDISS-8755 (2021).

55. G. Gerhardt, R. N. Adams, Determination of Diffusion Coefficients by Flow Injection Analysis. Anal Chem 54, 2618–2620 (1982).

56. D. Meyer, A. Hagemann, S. Kruss, Kinetic Requirements for Spatiotemporal Chemical Imaging with Fluorescent Nanosensors. ACS Nano 11, 4017–4027 (2017).

57. S. S. Nalige, P. Galonska, P. Kelich, L. Sistemich, C. Herrmann, L. Vukovic, S. Kruss, M. Havenith, Fluorescence changes in carbon nanotube sensors correlate with THz absorption of hydration. Nature Communications 2024 15:1 15, 1–8 (2024).

58. K. Ikeda, J. M. Bekkers, Counting the number of releasable synaptic vesicles in a presynaptic terminal. Proc Natl Acad Sci U S A 106, 2945–2950 (2009).

