## Supplementary Materials for "Rapid catecholamine trafficking regulates neutrophil functions and neutrophil-platelet interactions"

**The PDF file includes:**

Materials and Methods
Supplementary Text
Figs. S1 to S13
Tables S1 to S6
References (54-58)

### Materials and Methods

Isolation of human neutrophils from whole blood

This study was approved by the Ethics Committee Westfalen-Lippe (approval number 2021-657-f-S). Before donating blood, fully informed consent of each donor was obtained.

Neutrophils were isolated from whole blood of healthy donors by the density gradient separation method. Briefly, fresh blood was collected into EDTA blood collection tubes (Sarstedt, Germany). Whole blood was layered on Histopaque 1119 (Sigma-Aldrich, United States) and centrifuged at 1100 x g, for 21 min at room temperature with breaks off. The neutrophil layers (the third layer mainly containing density gradient and the fourth layer mainly containing neutrophils) were collected and diluted in Hank's Balanced Salt Solution (HBSS, Sigma-Aldrich, United States) without $Ca^{2+}$ and $Mg^{2+}$. The cell suspension was centrifuged at 400 x g, for 10 min at room temperature. The supernatant was discarded, and the cell pellet was resuspended in HBSS, layered on top of a five-step Percoll gradient (GE Healthcare, United States), and centrifuged at 1100 x g for 21 min at room temperature with breaks off. The neutrophils were collected from the fourth layer and a bit from the layers above and below the fourth. The neutrophils were diluted in HBSS and centrifuged at 400 x g, for 10 min at room temperature. The pelleted neutrophils were resuspended in RPMI 1640 cell medium (Gibco, United States) containing 10 mM HEPES (Sigma-Aldrich, United States), and 0.5 % fetal calf serum (FCS, heat-inactivated at 56 °C for 30 min, Gibco, United States) as needed.

For PCR and ELISA assays as well as FFN uptake experiments, neutrophils were isolated from whole blood using the EasySep™ Direct Human Neutrophil Isolation Kit (Stemcell Technologies, Canada) according to the manufacturer's instructions. The isolated neutrophil suspension was pelleted by centrifugation at 400 x g, for 10 min at room temperature and the pellet was taken up in RPMI 1640 cell medium containing 10 mM HEPES, and 0.5 % fetal calf serum (FCS, heat-inactivated at 56 °C for 30 min as needed. We confirmed the purity of neutrophils by a cytospin assay and staining using the Panoptical Fast Staining Kit (Carl Roth, Germany). We maintained neutrophil purity > 95 % of total isolated cells (without erythrocytes). Neutrophils were used for experiments immediately after isolation.

Immunofluorescence staining

1-2 x $10^5$ cells in 500 µL RPMI 1640 with 10 mM HEPES, 0.5 % FCS were seeded on round glass coverslips and left to adhere for 30-180 min at 37 °C, 5 % $CO_2$.

The adhered neutrophils were fixed by adding paraformaldehyde (PFA, biotium, United States or Morphisto, Germany) at a final concentration of 2 % for 15 min at room temperature. Cells were washed twice with PBS (Sigma-Aldrich, United States) and then permeabilized using PBS with 0.1 % Triton X 100 (Sigma-Aldrich, United States) for 10 min at room temperature. After two further washing steps with PBS, neutrophils were blocked with PBS containing 0.5-10 % bovine serum albumin (BSA, Capricorn Scientific, Germany) for 40 min at room temperature. Fixed cells were incubated with primary antibodies (Table S1) in PBS containing 0.5-1 % BSA overnight at 4 °C. Cells were washed three times with PBS and subsequently incubated with secondary antibody (Table S1) in PBS containing 0.5-1 % BSA for 1 h at 37 °C. After three further washing steps, the samples were stained with Hoechst 33342 (Invitrogen, United States) for 15 min and mounted using fluorescence mounting medium (Agilent, United States). The stained neutrophils were imaged with an Eclipse Ti2 microscope (Nikon Instruments, Japan) equipped with an Orca Fusion BT camera (Hamamatsu Photonics, Japan), an Axiovert 200 microscope (Zeiss, Germany) with a CoolSNAP ES camera (Photometrics, United States) or an LSM800 Airyscan confocal microscope (Zeiss, Germany).

For colocalization immunofluorescence staining of vesicular monoamine transporter 2 (VMAT2) and myeloperoxidase (MPO), the adhered cells were carefully washed twice with warm PBS and fixed with ice-cold methanol (VWR Chemicals, United States) for 10 min at 4 °C. The cells were washed twice with PBS and then blocked for 3 h with 1 % BSA (Carl Roth, Germany) in PBS at room temperature. The antibodies against VMAT2 and MPO as well as their isotypes (Table S1) were incubated with the cells overnight at 4 °C in 1 % BSA solution. The following day, the cells were washed twice with PBS. Subsequently, the second antibodies Alexa Fluor™ 555 and Alexa Fluor™ 635 (1:1000, Invitrogen, United States) (Table S1) were incubated with the cells for 1 h at room temperature in 1 % BSA solution. The cells were then washed twice for 10 min with PBS, twice with 0.1 % polyoxyethylene sorbitan monolaurate (Tween 20, Carl Roth, Germany) in PBS, and again twice in PBS for 10 min. Last 4′,6-diamidino-2-phenylindole (DAPI, Thermo Fisher Scientific, United States) was added for 10 min and rinsed twice afterward. Microscopy was performed using a Thunder Imagers DMi8 (Leica Microsystems, Germany). Excitation was conducted at a wavelength of 555 nm using a filter at 630 nm. Additionally, excitation was performed at 635 nm without a filter. For both cases, an excitation intensity of 70% and an exposure time of 40 ms were applied. A 63x oil immersion objective (HC PL APO 63x/1.40-0.60 OIL, Leica, Germany) was used.

qPCR

RNA was isolated from naïve neutrophils or neutrophils incubated with LPS (50 ng/mL, Sigma-Aldrich, United States) for 2 h at 37 °C, 5 % $CO_2$. The Monarch Total RNA Miniprep Kit (New England Biolabs, United States) was used following the manufacturer’s instructions for use with leukocytes. The RNA was eluted in 31 µL water and the RNA concentration was determined using a DS-11 spectrophotometer (DeNovix, United States). The isolated RNA was stored at -70 °C until further use. RNA was converted into complementary DNA (cDNA) by reverse transcription using the SuperScript IV First-Strand Synthesis System (Invitrogen, United States) and stored at -20 °C until further use. qPCR of synthesized cDNA was performed using the 5x HOT FIREPol EvaGreen qPCR Mix Plus (ROX) (Solis BioDyne, Estonia). All primers (Table S2Table S) were generated by Microsynth AG (Switzerland) and diluted to 8 µM with nuclease-free water.

For each sample, 4 µL 5x HOT FIREPol EvaGreen qPCR Mix Plus (ROX), 0.5 µL of both the forward and backward primers, and 10 µL of nuclease-free water were added to 5 µL of cDNA in the qPCR plate. The samples were mixed by pipetting and centrifuged shortly for one minute at 1000 rpm. The qPCR reaction was run and measured on a qTOWER 2.2 quantitative real-time PCR thermal cycler with qPCRsoft 3.1 software (Analytik Jena, Germany) according to the thermal cycling profile described in Table S3. After the end of the qPCR reaction, a melting curve from 60 °C to 95 °C was generated.

Catecholamine, HVA, and VMA detection by ELISA

For cell lysis, isolated neutrophils were washed with cold PBS and then resuspended in 150 µL cold PBS. The cell suspension was tip-sonicated with a model 120 sonic dismembrator (Fisherbrand, United States) at 70 % amplitude, 10 s on / 20 s off cycle, for 3.5 min on ice. The sample was then frozen in liquid nitrogen and thawed in an ultrasonic bath (Bransonic 221, Branson Ultrasonics, United States) three times. Cell debris was pelleted by centrifugation at 14,000 x g, 4 °C, for 10 min and the lysate was further cleaned by centrifugation through a nylon micro-centrifuge filter (0.2 µm, Ciro, United States) at 1000 x g, 4 °C, for 2 min. Neutrophils were

incubated with latrunculin A (1 µM, Tocris Bioscience, United Kingdom) for 30 min and afterwards stimulated with N-Formylmethionyl-leucyl-phenylalanine (fMLP, 1 µM, Merck, Germany) for 5 min at 37 °C. Cells were then centrifuged at 4602 x g for 2 min, the supernatant was treated with 1 x Halt™ Protease Inhibitor (Thermo Scientific, United States) while the cells were lysed for later analysis.
ELISAs for dopamine (Abcam, United Kingdom), norepinephrine (Abcam, United Kingdom), epinephrine (Abcam, United Kingdom), HVA (Abbexa, United Kingdom), and VMA (Abbexa, United Kingdom) were performed according to the manufacturer's instructions. The measurements were performed using a Hidex Sense 425-301 (Hidex, Finland) microplate reader with a readout of ng/mL or pg/mL. The cell number per sample and sample volume were used to calculate the concentration per cell. The volume of a neutrophil was assumed as 1 pL.

Fluorescence Microscopy of FFN Uptake by Neutrophils
300,000 cells were seeded in a 3.5 cm cell culture dish and incubated for 30 min to ensure that they sedimented and adhered to the bottom of the dish. After sedimentation, the cells were washed once with HBSS with 2 mM calcium chloride ($CaCl_2$, Carl Roth, Germany). HBSS was then added to the cells in the cell culture dish. Microscopy was performed using a Thunder Imagers DMi8 (Leica Microsystems, Germany). Excitation was performed at 475 nm without filter with an intensity of 70 % and an exposure time of 40 ms. A 63x oil immersion objective (HC PL APO 63x/1,40-0,60 OIL, Leica, Germany) was used. Before starting the measurement, an image was taken in bright field mode, focusing on the vesicles, which appeared as black structures. Then the measurement was started and fluorescent false neurotransmitters 511 (FFN511, Abcam, United States) dissolved in DMSO (Sigma-Aldrich, United States) were added simultaneously at a concentration of 2.5 µM and the uptake was measured for 10 min. The images were obtained from Leica LAS X Thunder software and analyzed with ImageJ/Fiji.

Primary murine neuron culture and immunofluorescence staining
Primary ventral midbrain neurons were isolated from C57BL/6N mice. They were grown on an astrocyte feeder layer isolated from C57BL/6N cortex for 2-6 weeks before use in immunofluorescence experiments. Samples were prepared, fixed, and labeled as published previously. *(54)*

Fluorescence microscopy of FFN exocytosis from neutrophils
Neutrophils were adhered on glass chamber slides at $10^6$/mL concentration in HBSS (with $Ca^{2+}$ and $Mg^{2+}$). FFN511 was dissolved in DMSO and added to the cell suspension with the final concentration of 5 µM. Cells were incubated at 37 °C, 5 % $CO_2$, for 5 min, then media was aspirated and cells were washed 3 times with HBSS and immediately used for experiments. For FFN102 (Abcam, USA), 10 µM concentration and 30 min incubation time was used. Cells were then triggered by adding 100 µM serotonin (Thermo Fisher Scientific, United States) and the exocytosis was imaged using an Olympus IX83 microscope (Japan) coupled with a CoolLed pE-4000 illumination system. The FFNs were excited with a 488 nm LED at 500 ms exposure time. Image sequences were acquired before, during, and after the stimulant was added with a pipette to the chamber slide. The images were obtained from Olympus cellSense software and analyzed with ImageJ.

Synthesis of nanosensors

(6,5)-chirality enriched SWCNTs (carbon ≤ 95 %, ≥ 93 % carbon as SWCNT, Signis® SG65i, 0.7 – 0.9 nm diameter (Sigma-Aldrich, United States) and $(GT)_{10}$ oligonucleotide (5'-GTGTGTGTGTGTGTGTGTGT-3 synthesized by Sigma-Aldrich, United States) were mixed in phosphate saline buffer (Gibco, United States) with final concentrations of 0.5 mg/mL and 50 µM respectively. The dispersion was tip-sonicated with a Model 120 Sonic Dismembrator (Fisherbrand, United States) at 30 % amplitude for 20 min. The nanosensor suspension was centrifuged at 16,000 × g for 30 min twice. After each centrifugation, the pellet was discarded and the supernatant was collected for further investigation. The nanosensor suspension was stable at room temperature for several months.

Nanosensor characterization

Absorption spectra of the nanosensors were collected with a UV–vis–nIR spectrometer (JASCO V-670, Spectra Manager Software). They were diluted 1:100 in PBS to measure nIR absorbance. The concentration was then calculated by integrating the area under the curve belonging to (6,5) SWCNTs peak and taking the extinction coefficient with an estimated length of 600 nm. For all catecholamine-sensing experiments with neutrophils, the applied concentration was 4 nM. For acquiring the fluorescence emission spectra of the nanosensors in suspension the concentration was 2 nM. For fluorescence spectra the nanosensor suspension in HBSS (2 nM) was excited with a 561 nm laser coupled to an Olympus IX73 microscope at 1 s integration time. The emission spectra from 800 nm to 1300 nm were acquired by an Andor iDus InGaAs 491 array NIR detector attached to a Shamrock 193i spectrograph (Andor Technology Ltd., Northern Ireland).

Fluorescence Microscopy of Neutrophils Adhered on Nanosensors

Nanosensors were immobilized on glass surfaces by incubating the surface with 4 nM nanosensors suspension overnight at 4° C. Before measurements, the surfaces were washed with HBSS three times and HBSS (with 2 mM $Ca^{2+}$ and $Mg^{2+}$) was added to the surface. Neutrophils at a concentration of $10^6$/mL in HBSS (with 2 mM $Ca^{2+}$ and $Mg^{2+}$) were seeded onto the nanosensor immobilized glass bottom petri dishes or slides. The neutrophils were let to adhere for 5 min and the medium was aspirated to remove loosely adhered cells. Fresh medium was added to the cells and image acquisition was started. After typically 10 s, the stimulant was added to the media while image acquisition continued.

For imaging, we employed an Olympus BX53 or IX73 microscope, a 561 nm laser (Cobolt Jive™ laser, Cobolt AB, Sweden), and two cameras. An Andor Zyla 5.5 sCMOS camera, (Andor Technology Ltd., UK) for visible fluorescence and NIR InGaAs (Cheetah-640-TE-1, Xenics, Belgium) camera for nIR fluorescence. Images were obtained with a 100x objective lens (UPLSAPO100XS, Olympus, Japan) and recorded with down to 100 ms exposure time and up to 15 frames per second. For visible fluorescence, a xCite 120Q fluorescence lamp and an EGFP excitation filter were used. The emission was filtered by a 650 nm short-pass filter, a 561 nm notch filter, and a 525/50 nm bandpass filter. For nIR fluorescence the 561 nm laser was used typically at 100 mW power. Images were analyzed with ImageJ. The drift was corrected by subtracting a linear line.

Calcium Flux Measurement

Neutrophils at $10^6$/mL concentration in RPMI 1640 (supplemented with 10 % FBS) were incubated with Fluo-4, AM (1 µM, Thermo Fisher Scientific, United States) at room temperature for 40 min. The cells were washed once and resuspended in HBSS (with $Ca^{2+}$ and $Mg^{2+}$). The cell suspension was added to glass-bottom, black 96-well plates. A CLARIOstar Plus plate reader was employed. Wells were scanned with (excitation 480-14 nm/emission 530-30) 691 times at 0.41 s intervals. At t = 50 s and t = 250 s, an autoinjector pumped 10 µL of stimulant with 430 µL/s speed into the well.

CA release, diffusion, and sensor response simulation

A numerical simulation was performed (in 2D) to model the release of CA (with the diffusion constant of dopamine/DA), the subsequent diffusion, and the binding dynamics of the DA molecules to nanosensors. The simulation can be divided into two main processes: A simulation of the release and diffusion of DA, and a simulation of the binding of DA molecules to the (nano)sensors based on the spatiotemporal concentration profile. Both processes were simulated with numerical methods to ensure accuracy and efficiency:

A numerical simulation of the diffusion equation (Fick's second law) was chosen to model the diffusion. For this purpose, a Runge-Kutta method was applied, which is particularly suitable for approximating time-dependent partial differential equations such as the diffusion equation. To describe the experimental situation, a 2-dimensional geometry of the diffusion was assumed. The structure of substrate, sensors, and cells within a thin liquid layer only allows diffusion in two dimensions. The diffusion area was divided into pixels, for each pixel, and for each time step, the dopamine concentration was calculated based on the concentration of the surrounding pixels in the previous time step. The size of the time steps was chosen sufficiently small to ensure high accuracy and stability of the simulation.

To test the reliability of the simulation, a simple geometry was considered after a instantaneous release from a point source, for which the diffusion equation can also be solved analytically in two dimensions (Figure S7a). A comparison of the simulation and the analytical solution shows only minimal deviations. The simulation only deviates from the analytical solution due to the selected boundary conditions for very long diffusion times and low concentrations that are no longer relevant for sensory analysis.

The release of DA can be incorporated into the simulation in various ways. A release at different times and at different locations is possible, but also a single release event. The release can either be simulated as coming from a point-like source or evenly distributed over the complete surface of a cell.

The binding process was modeled probabilistically, accounting for the stochastic nature of molecular interactions. Binding events are governed by rate constants for binding and unbinding reactions. At each time step, the simulation checks each sensor for potential binding and unbinding events based on the local concentration of molecules and the number of available binding sites. This method ensures a realistic representation of the dynamic equilibrium between free and bound molecules. Every binding or unbinding event changes the local concentration in the area of the sensor. Since only one binding or unbinding event is considered per binding site and time step, the size of the time steps had to be chosen sufficiently small so that the probability of multiple binding and unbinding events at a single binding site can be neglected.

For the simulation, a series of parameters had to be tailored to fit the biological conditions, as given in Table S4 and Table S5. Several runs of the simulation with different changes in the parameters were performed. Each run of the simulation leads to DA concentration mappings and bound DA molecule number mappings for each simulated time step. For analysis, two central outputs were evaluated for each run of the simulation: The number of free DA molecules in the area of the cell (a circle with radius 10 µm in the center of the simulated area) and the number of sensor-bound DA molecules in the same area. As the bound DA molecules on the SWCNT are the cause of the fluorescence increase, we can compare the simulated number of bound DA molecules with the experimental results of the fluorescence intensity.

Out of all the five considered runs of the simulation, run 1 fits the experimental results (Fig. 3D) qualitatively best. The number of bound DA molecules increases over the course of ~30 s, before the increase decelerates and binding and unbinding come close to reaching an equilibrium. After the last release event, an unbinding due to the sensor kinetics given by $k_{off}$ can be observed.
After investigating other simulated parameters, a scenario of a release separated into several release events over the course of several 10 s to 1 min combined with sensor kinetics described by an on rate in the order of $10^6$ $M^{-1}s^{-1}$ and an off rate in the order of 0.1 $s^{-1}$ describes the observed results best.
The observed slow increase in the fluorescence signal cannot be caused by a single release event at one time (run 2, run 4). Due to the diffusion, already a few seconds after the release event, the DA concentration would have decreased below the detection level of the nanosensors. A mathematically possible exception to this is the case where diffusion is reduced by several orders of magnitude compared to the literature values *(55)* and the on-rate is also reduced by several orders of magnitude (run 5). In this case, however, the ratio between on rate and off rate would no longer agree with the $K_d$ values of the DA nanosensors known from literature *(24,28)* and this option can be rejected.
Compared to even slower off-rates (run 3), the parameters chosen in run 1 fit the experimental results better, considering the slower increase after ~30 s and the speed of the signal decrease after the last release event. Additionally, the ratio between on rate and off rate fits best with the $K_d$ values of the nanosensors known from literature. *(24,28)*

Isolation of human platelets from whole blood
Platelets were isolated from whole blood of healthy donors. This study was approved by the Ethics Committee Westfalen-Lippe (approval number 2021-657-f-S). Before donating blood, fully informed consent of each donor was obtained.

Whole blood was collected into citrate 9NC blood collection tubes (Sarstedt, Germany) and centrifuged at 259 x g, for 30 min. Platelet-rich plasma (PRP) was separated and centrifuged at 259 x g, for 10 min. PGE1 at a final concentration of 6.6 µM was added to the PRP and centrifuged at 10,000 x g for 30 s. The pelleted platelets were carefully resuspended in Tyrode’s solution (pH 6.2) supplemented with PGE1 at a final concentration of 5 µM. Then, the cell suspension was centrifuged again at 10,000 x g for 30 s and resuspended in Tyrode’s solution (pH 6.2) with PGE1 as described before. Finally, the platelets were centrifuged at 10,000 x g for 15 s, resuspended in Tyrode’s solution (pH 7.4), and kept at 37 °C before starting experiments.

5-HT receptor inhibition
Nanosensors were immobilized on glass surfaces by incubating the surface with 4 nM nanosensor suspension overnight at 4° C. Neutrophils at $10^6$/mL concentration in RMPI 1640 were seeded onto the nanosensor-coated glass surface and incubated with or without 100 µM ketanserin (+)-tartrate (ketanserin, Sigma-Aldrich United States) and subsequently washed with buffer.

Stimulation with either 100 µM 5-HT or 1 µM 2-[(3-chlorophenyl)methoxy]-6-(1-piperazinyl)-pyrazine (CP809, Tocris Bioscience, United Kingdom) was performed 10 s after the start of imaging. Images were obtained with a 100x objective lens (UPLSAPO100XS, Olympus, Japan) and recorded with 100 ms exposure time and at 5 frames per second. For nIR fluorescence imaging the 561 nm laser was used at 100 mW power. Images were analyzed with ImageJ. Drift was corrected by subtracting a linear line.

Neutrophil-platelet interaction

Nanosensors were immobilized on glass surfaces by incubating the surface with 4 nM nanosensors suspension overnight at 4° C. For control measurements, thrombin (1 Unit/mL, Sigma-Aldrich, United States) was added onto blank nanosensors or onto nanosensors with adhered platelets. For neutrophil measurements, neutrophils at $10^6$/mL in HBSS were seeded on a nanosensor-coated glass surface and incubated for 5 min before platelets were added and incubated for 5 min. The measurement was started, and thrombin (1 Unit/mL) was added after 10 s. Images were obtained with a 100x objective lens (UPLSAPO100XS, Olympus, Japan) and recorded with 200 ms exposure time and at 5 frames per second rate. For nIR fluorescence 561 nm laser was used at 120 mW power. Images were analyzed with ImageJ. The drift was corrected by subtracting a linear line.

For serotonin receptor inhibition neutrophils and platelets were seeded onto the nanosensors and treated with 100 µM ketanserin for 10 min prior to stimulation. The cells were rinsed with HBSS. The measurement was started and thrombin (0,01 Unit/mL) was added after 10 s. Images were obtained with a 100x objective lens (UPLSAPO100XS, Olympus, Japan) and recorded with 200 ms exposure time and at 5 frames per second rate. For nIR fluorescence 561 nm laser was used at 120 mW power. Images were analyzed with ImageJ.

NETosis assay

Neutrophils were seeded on glass bottom 96-well plates (10,000 per well), activated with PMA (100 nM, Sigma-Aldrich, United States), and simultaneously incubated with dopamine hydrochloride, DL-norepinephrine hydrochloride, (±)-epinephrine hydrochloride or serotonin hydrochloride (all Sigma-Aldrich, United States) at 37 °C, 5 % $CO_2$. After a 3 h incubation time, the cells were fixed with 2 % PFA to stop NET formation and stored overnight at 4 °C. Cells were then washed once and the chromatin was stained with Hoechst 33342 at room temperature. Neutrophils were imaged with an Eclipse Ti2 microscope (Nikon Instruments) equipped with an Orca Fusion BT camera (Hamamatsu Photonics) or an Axiovert 200 microscope (Zeiss) with a CoolSNAP ES camera (Photometrics). Eight images from random regions were collected for each well. For all experiments, the number of decondensed nuclei and the total cell count were quantified with ImageJ.

Aggregometry

Aggregometry experiments were performed using an eight-channel aggregometer (PAP-8E, möLab, Germany). Before the start of the experiment, $MgCl_2$ and $CaCl_2$ (both Sigma-Aldrich, United States) were added to the platelets to achieve a final concentration of 1 mM and 2 mM, respectively. The platelets were incubated with the catecholamines (dopamine hydrochloride, DL-norepinephrine hydrochloride, (±)-epinephrine hydrochloride, all Sigma-Aldrich, United States) for approximately 2 min before stimulation with thrombin (1.2 U/mL, Sigma-Aldrich, United States) and the platelet response was measured throughout. The measurements were continued for

30 min or until the aggregation had reached a plateau. The magnitude of aggregation was determined by finding the maximum aggregation throughout the measurement. The aggregation speed was determined by performing a Michaelis-Menten least squares curve fit of the aggregation data generated in the first 20 min after the stimulation of aggregation and is presented as the $K_m$ value.

Statistics

Statistical tests were performed with single-cell data when available. Data was checked for normality using Shapiro-Wilk tests and consequent statistical tests were chosen accordingly. For n numbers (independent donors) and N numbers (individual cells), see Table S6.

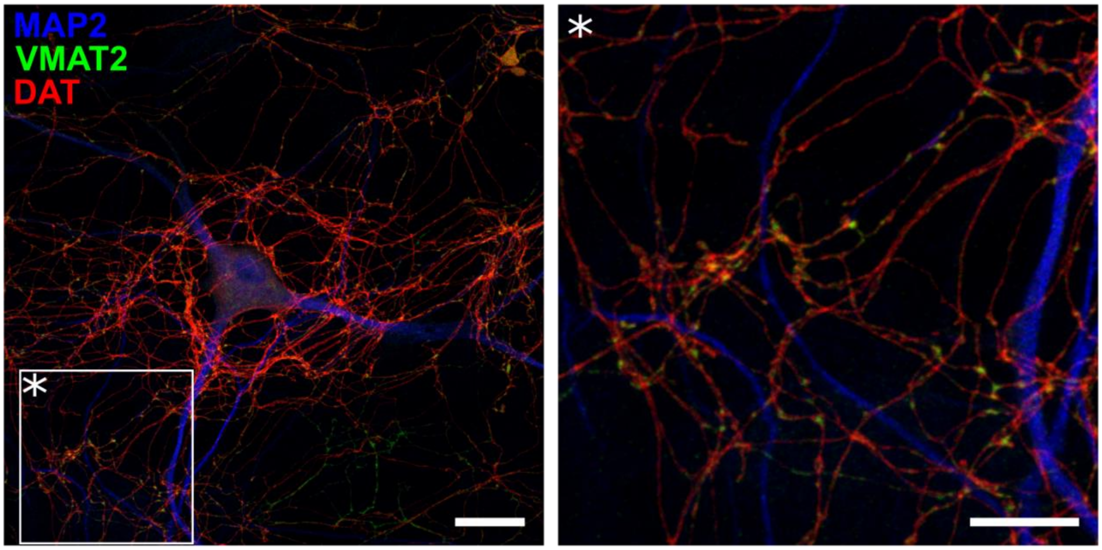


**Fig. S1.** Primary mouse ventral midbrain neurons cultured on astrocytes and stained for microtubule-associated protein 2 (MAP2, blue), vesicular monoamine transporter 2 (VMAT2, green) and dopamine transporter (DAT, red). Scale bar is 20 µm in the overview (left) and 10 µm in the detailed view (right).

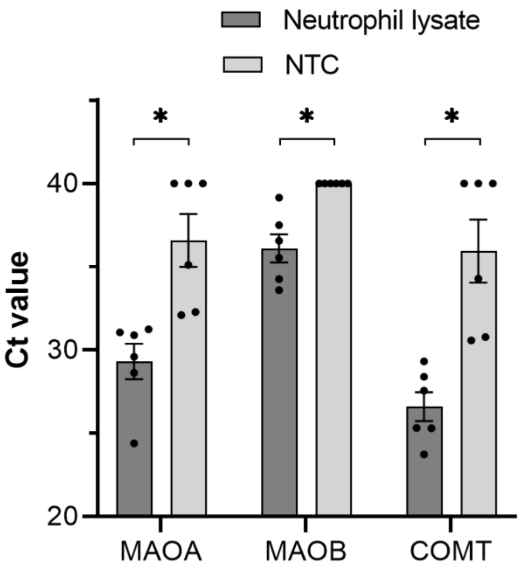


**Fig. S2.** *MAOA, MAOB, and COMT are expressed at the RNA level as determined by qPCR (n = 6, mean ± SEM, Wilcoxon matched-pairs signed rank test, NTC: no template control).*

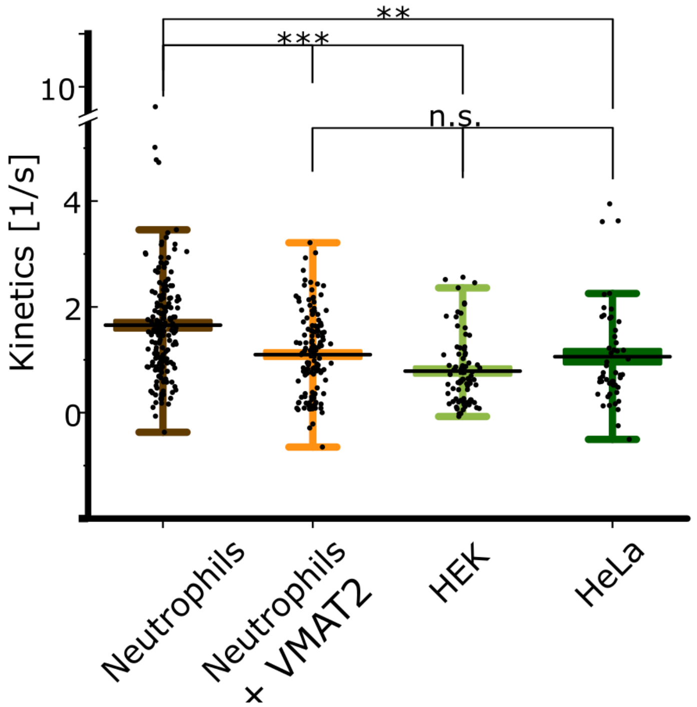


**Fig. S3.** Uptake kinetics of 2.5 nM FFNs of control neutrophils, inhibited neutrophils (+ VMAT2 inhibitor), HEK, and HeLa cells (n = 5, mean ± SEM, one-way ANOVA, p-value ** < 0.01, *** < 0.001).

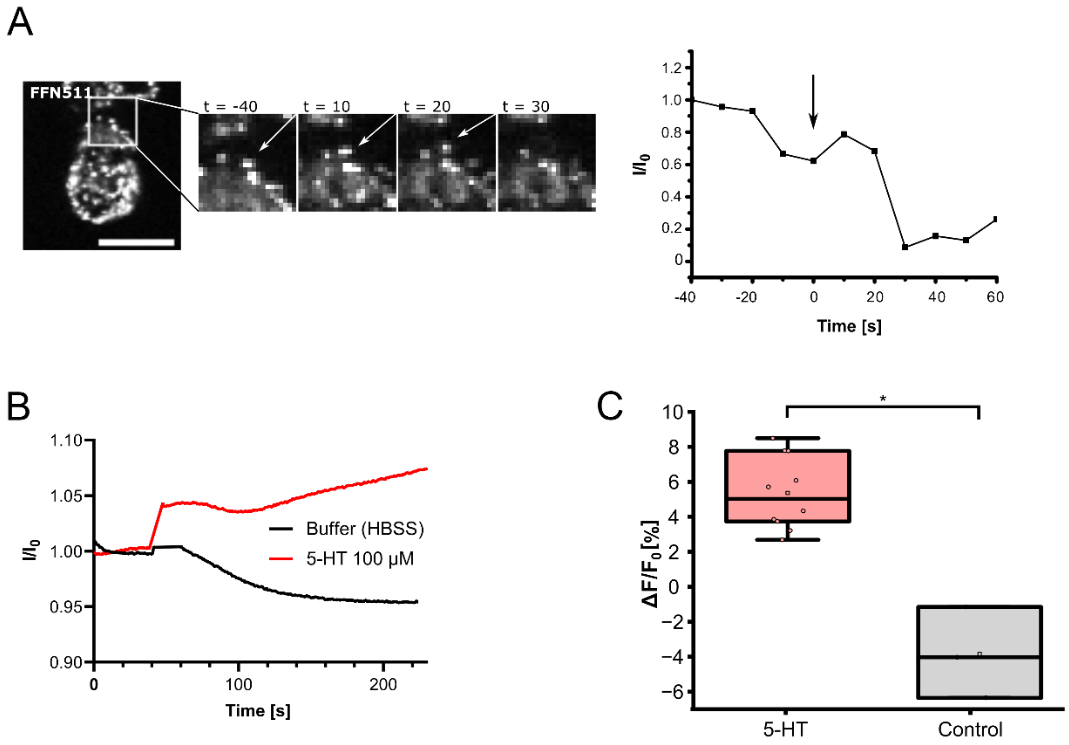


**Fig. S4.** (**A**) Tracking of single FFN511 vesicles in a neutrophil before, during, and after stimulation with 100 µM serotonin, over time. Vesicle trafficking toward cell membrane, fusion with membrane, and extracellular release of FFN511 are observed and analyzed and steps in the intensity trace (right) indicate exocytosis. Scale bar is 10 µm. (**B**) Fluorescence intensity changes of the extracellular region around an exemplary cell overtime when cells are stimulated with serotonin (5-HT) compared to the control cell which is stimulated with HBSS (cell incubated with FFN102). The increase of extracellular fluorescence indicates exocytosis. (**C**) Endpoint measurements of fluorescence intensity of the extracellular region around the cells over time when cells are stimulated with serotonin (5-HT) compared to control cells stimulated with HBSS (cells incubated with FFN102) ($n = 1$, $N_{5\text{-HT}} = 10$, $N_{Control} = 3$, mean ± SEM, paired t-test, p-value * $< 0.05$).

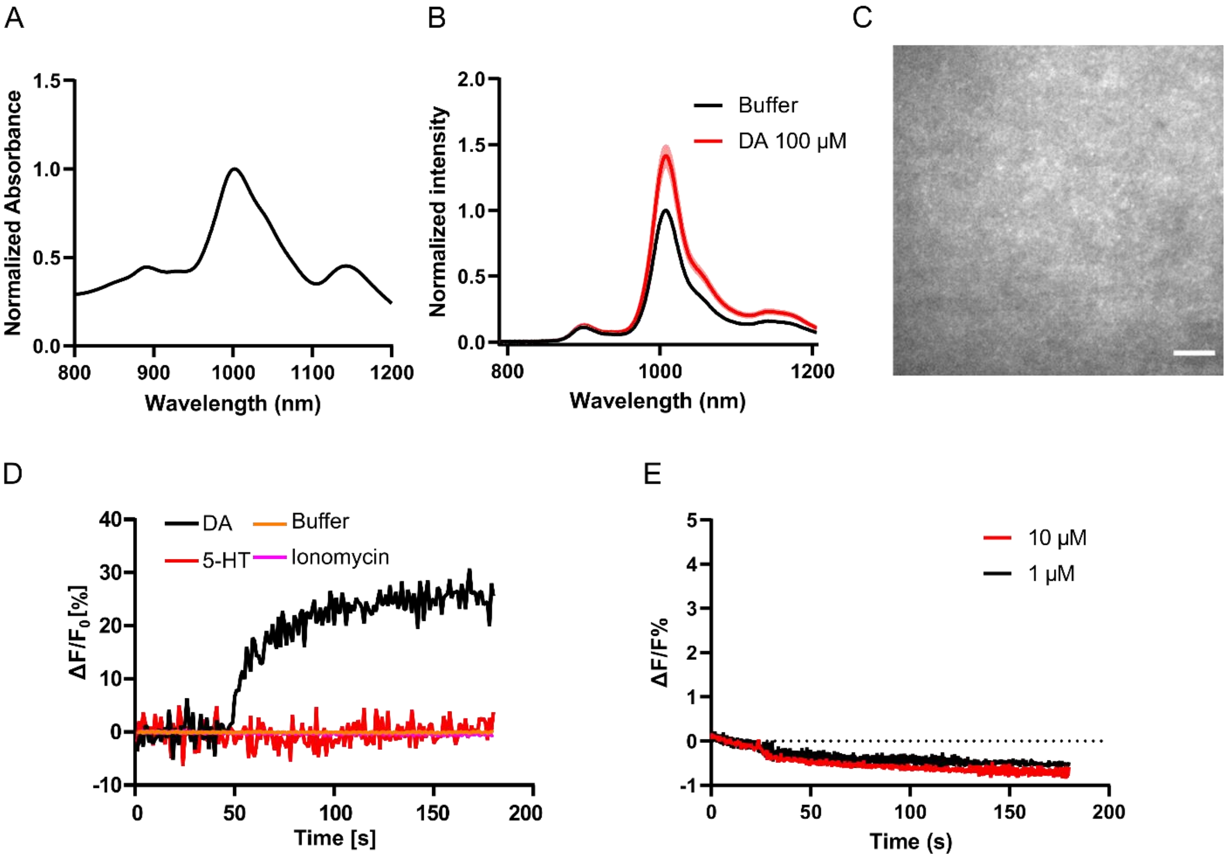


**Fig. S5.** (**A**) nIR absorbance spectrum of nanosensor suspension. (**B**) Normalized fluorescence emission spectra of nanosensor in suspension before and after dopamine (DA) addition in PBS. Shades = SD (n = 3). (**C**) Nanosensor surface coverage imaged with a nIR camera. Scale bar is 10 µm. (**D**) The fluorescence response of nanosensors to dopamine, serotonin, ionomycin (blank nanosensors), and buffer (neutrophil-adhered on nanosensors) over time. (**E**) Fluorescence signal of nanosensor exposed to $H_2O_2$ in PBS (t = 25 s) does not increase and the decrease is most likely a z-drift of the focus.

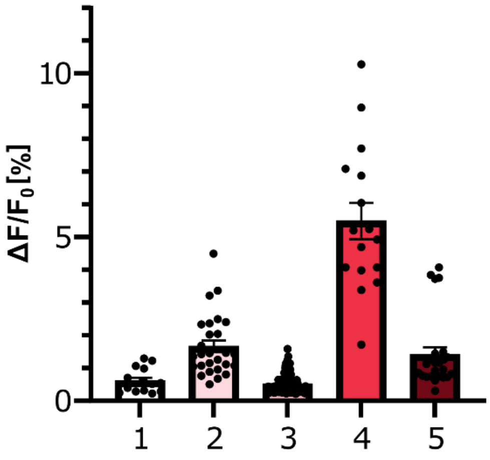


**Fig. S6.** The maximum fluorescence response of nanosensors after release of CA from neutrophils stimulated by serotonin. Each bar represents an independent experiment from a different blood donor and each dot is a single neutrophil (mean ± SEM).The data show the heterogeneity between individual cells, which could also be attributed to different adherence/distance to the nanosensor layer.

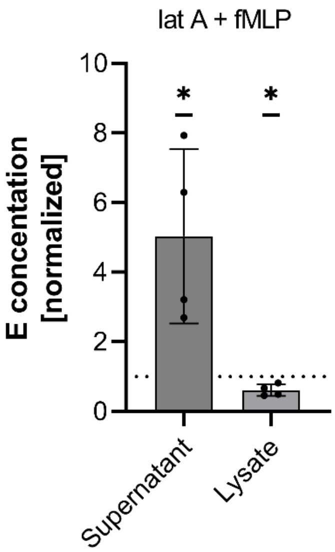


**Fig. S7** Normalized epinephrine concentration in neutrophil supernatant and lysate after pre-treatment with latrunculin A and stimulation with fMLP as determined by ELISA (n = 4, mean ± SEM, one sample t-test: p-value * < 0.05).

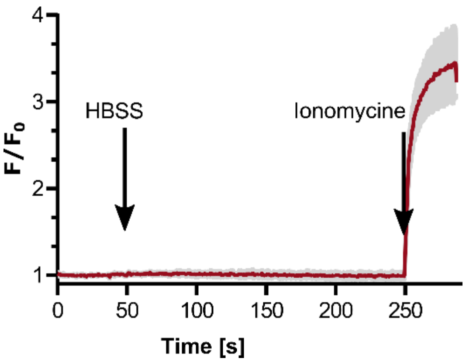


**Fig. S8.** Fluorescence signal of exemplary neutrophils incubated with Fluo-4 AM, $Ca^{2+}$ indicator, over time when HBSS (negative control) is added at t = 50 s and ionomycin ($Ca^{2+}$ ionophore, positive control, 5 µM) is added at t = 250 s.

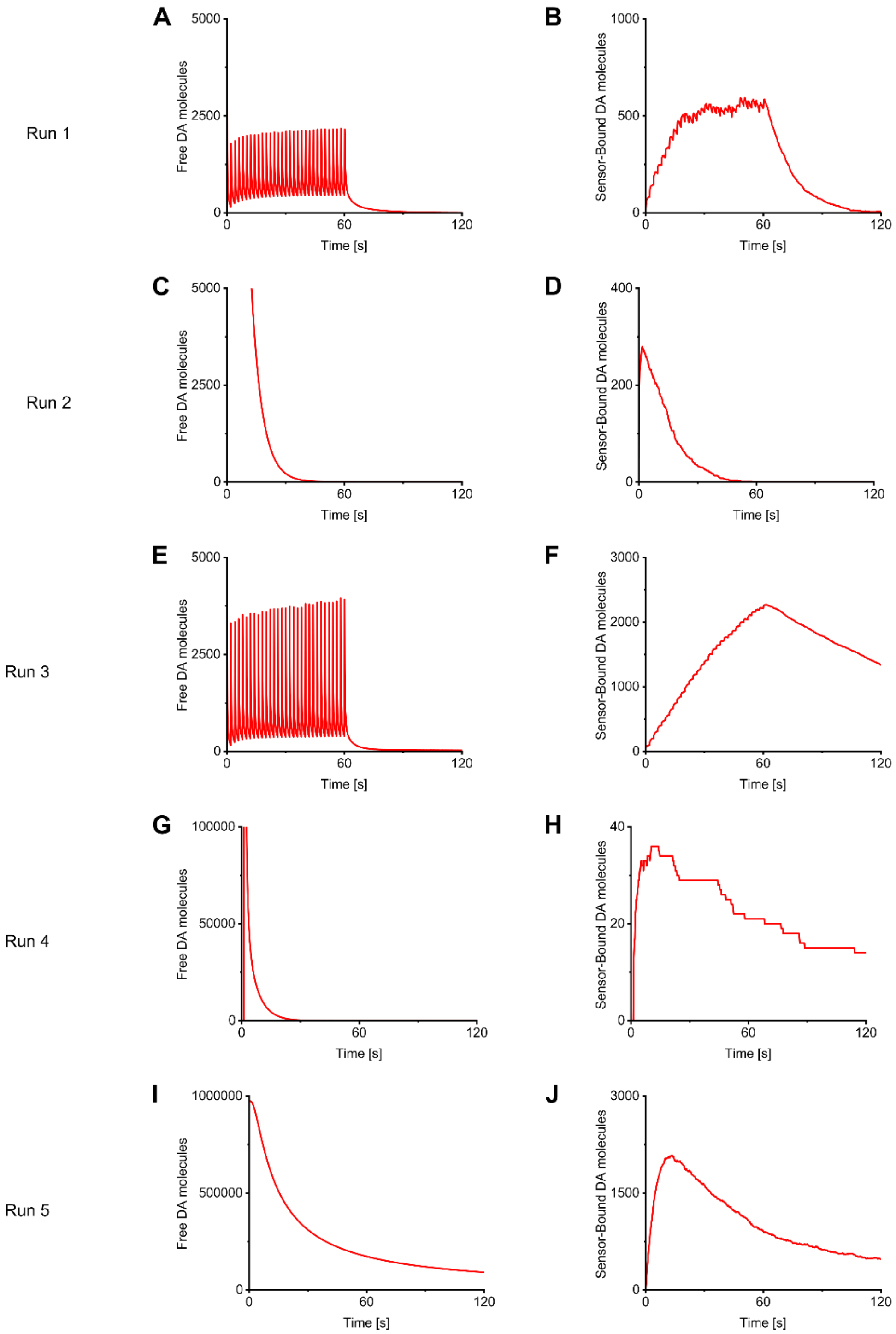


**Fig. S9.** DA release, diffusion, and sensor response simulation: A, C, E, G, I: Free DA molecules under the area of the simulated cell in the center of the simulation area. B, D, F, H, J: Number of sensor-bound DA molecules (=sensor response) in the area of the simulated cell in the center of the simulation area. Note that only the sensor response can be measured experimentally. Data for five different preliminary runs of the simulation according to Table S5.

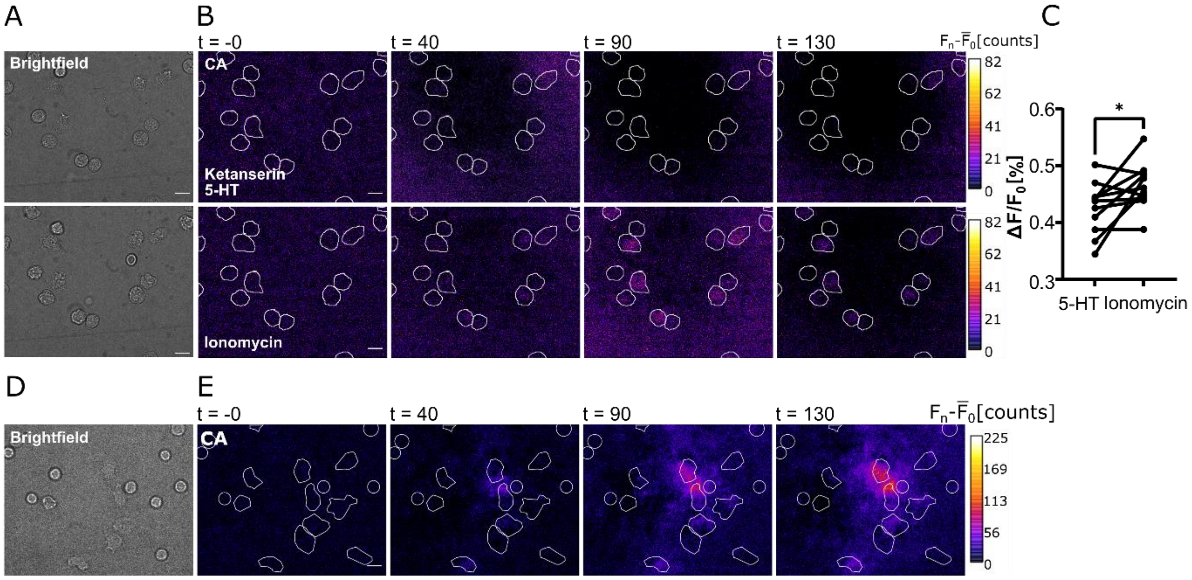


**Fig. S10.** (A) Brightfield image and (B) nIR image sequences of neutrophils adhered on a nanosensor surface. Background is measured 10 s before addition of serotonin ($\overline{F}$). Neutrophils are first incubated with ketanserin (100 µM) and stimulated with serotonin (100 µM) at t = 0 (top). The same cells were then stimulated with 5 µM ionomycin (bottom). (C) Maximum fluorescence signal after stimulation of exemplary cells in section (A, B) (n =1, paired t-test: p-value * < 0.05). (D) Brightfield image and (E) nIR image sequence of neutrophils adhered on nanosensor surface before and after stimulation with CP809 (1 µM). Background is measured 10 s before addition of serotonin ($\overline{F}$).

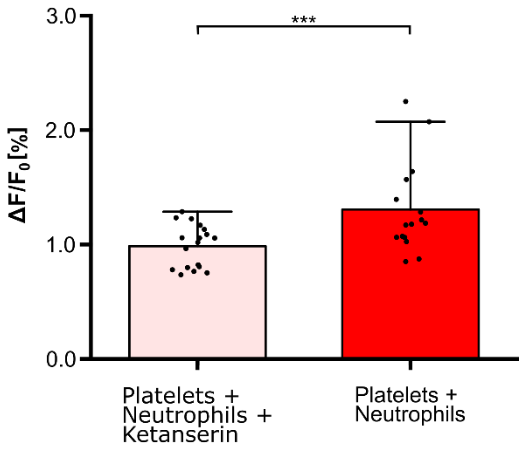


**Fig. S11.** The maximum fluorescence response of nanosensors after release of CA from neutrophils. Neutrophils were stimulated by platelets, which were stimulated with thrombin (n = 1, mean ± SEM, Mann-Whitney tests: p-value ** < 0.01, each data point corresponds to a single neutrophil).

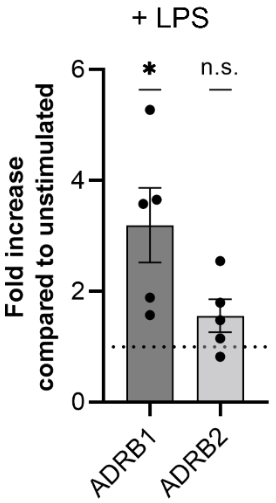


**Fig. S12** The expression of ADRB1 but not ADRB2 is increased after stimulation of neutrophils with 50 ng/mL LPS for two hours (n = 5, mean ± SEM, one-sample t-test).

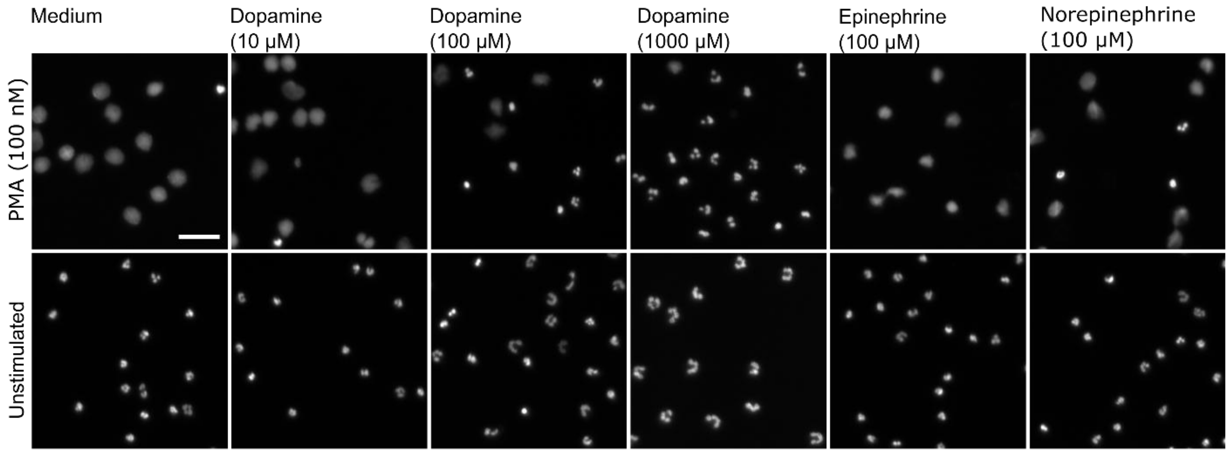


**Fig. S13** NETosis assay of neutrophils, stimulated with PMA (top) or unstimulated (bottom) and treated with different concentrations of catecholamines. The chromatin is stained with Hoechst. Scale bar is 40 µm.

**Table S1. Antibodies used for immunofluorescence staining**

| **primary antibodies** | **supplier** | **concentration** |
|---|---|---|
| anti-β1-adrenergic receptor polyclonal antibody – rabbit IgG | Invitrogen, PA1-049 | 1:200 |
| anti-β2-adrenergic receptor polyclonal antibody – rabbit IgG | Invitrogen, PA5-80323 | 1:1000 |
| anti-catechol-O-methyltransferase polyclonal antibody – rabbit IgG | Novus Biologicals, NBP3-03726 | 1:100 |
| anti-dopamine transporter polyclonal antibody – rabbit IgG | Proteintech, 22524-1-AP | 1:200 |
| anti-dopamine transporter antibody, culture supernatant – rat IgG2a κ | Sigma-Aldrich, MAB369 | 1:1000 |
| anti-monoamine oxidase A monoclonal antibody – rabbit IgG | Abcam, ab126751 | 1:40 |
| anti-monoamine oxidase B polyclonal antibody – rabbit IgG | Abcam, ab175136 | 1:100 |
| anti-microtubule-associated protein 2 polyclonal antibody – chicken IgY | Novus Biologicals, NB300-213 | 1:2000 |
| anti-myeloperoxidase monoclonal antibody – rat IgG2a | Abcam, 1055121-21 | 1:500 |
| anti-tyrosine hydroxylase polyclonal antibody – rabbit antiserum | Synaptic Systems, 213102 | 1:500 |
| anti-vesicular monoamine transporter 2 polyclonal antibody – rabbit IgG | Invitrogen, PA5-112713 | 1:200 |
| anti-vesicular monoamine transporter 2 polyclonal antibody – rabbit IgG | Proteintech, 20873-1-AP | 1:200 |
| anti-vesicular monoamine transporter 2 antibody – rabbit IgG | Frontier Institute, AB2571857 | 1:2000 |
| **isotypes** | | |
| rabbit IgG monoclonal antibody | Abcam, ab172730 | according to primary |
| rabbit IgG monoclonal antibody | Invitrogen, 08-6199 | undiluted |
| rat IgG2a monoclonal antibody | Invitrogen, 02-9688 | 1:500 |
| **secondary antibodies** | | |
| goat anti-rabbit IgG secondary antibody – Alexa Fluor™ 488 | Invitrogen, A-11034 | 1:500 |
| goat anti-mouse IgG secondary antibody – Alexa Fluor™ Plus 555 | Invitrogen, A32727 | 1:1000 |
| goat anti-rabbit IgG secondary antibody – Alexa Fluor™ 635 | Invitrogen, A-31576 | 1:1000 |

**Table S2. Primer sequences for qPCR**

| gene | direction | sequence |
|---|---|---|
| ADRB1 | forward | 5‘-CCGGGAACAGGAACACAC-3‘ |
| ADRB1 | reverse | 5‘-GAAAGCAAAAGGAAATATGTCTTGA-3‘ |
| ADRB2 | forward | 5‘-TTGCCTCTTCCATCGTGTCC-3‘ |
| ADRB2 | reverse | 5‘-CCACCTGGCTAAGGTTCTGG-3‘ |
| B2M | forward | 5‘-CCACTGAAAAAGATGAGTATGCCT-3‘ |
| B2M | reverse | 5‘-CCAATCCAAATGCGGCATCTTCA-3‘ |
| COMT | forward | 5‘-GGAGGCCATTGACACCTACTG-3‘ |
| COMT | reverse | 5‘-CGATCTTGCCTTTCTTGTCGC-3‘ |
| MAOA | forward | 5‘-AGGACTATCTGCTGCCAAAC-3‘ |
| MAOA | reverse | 5‘-AAGCTCCACCAACATCTACG-3‘ |
| MAOB | forward | 5‘-GCGGCATCTCAGGTATGGCA-3‘ |
| MAOB | reverse | 5‘-TCCAATCCTAGCTCCTTGGCT-3‘ |

**Table S3. Thermal cycling profile for qPCR**

| step | temperature | duration | cycles |
| --- | --- | --- | --- |
| initial denaturation | 95 °C | 15 min | 1 |
| denaturation | 95 °C | 15 s | 40 |
| annealing | 56 °C | 20 s | |
| elongation | 72 °C | 20 s | |
| melting curve | 60 – 95 °C, $\Delta T = 1$ °C | 10 s | 1 |

**Table S4. Parameters for simulation**

| **Parameter** | **Chosen Value** | **Source** |
|---|---|---|
| Simulation area: | 256 µm x 256 µm | |
| Resolution (Pixel Size): | 0.5 µm x 0.5 µm | |
| Simulated time | 120 s | According to the experimental conditions |
| Placement of cell in simulation area | 10 µm diameter, in the center of the | According to the experimental conditions |
| Density of SWCNT nanosensors | 0.5 µm x 0.5 µm per sensor (1 per pixel) | based on *(56)* |
| Binding sites per nanosensor | 30 | based on *(57)* |
| $k_{on}$ | V | |
| $k_{off}$ | Varies between the different runs of the simulators | |
| DA diffusion coefficient | Varies between the different runs of the simulators | |
| Geometry of DA release | Varies between the different runs of the simulators | |
| Amount of DA molecules released | Varies between the different runs of the simulators | |

**Table S5. Variation of parameters for different runs of the diffusion simulation**

| | $k_{on}$ | $k_{off}$ | **Diffusion coefficient** | **Geometry of DA release** | **Amount of DA molecules released** |
|---|---|---|---|---|---|
| **Run 1** | $10^6$ $M^{-1}s^{-1}$ (based on *(56)*) | 0.1 $s^{-1}$ (based on*(56)*) | D=6.05*$10^{-6}$ cm²/s (based on *(55)*) | Release from small vesicles at a random position on the edge of the cell, 30 release events every 2 s (own assumption, singular release events from different vesicles given e.g. in*(58)*) | 30000 DA molecules per release event (as in *(58)*) |
| **Run 2** | $10^4$ $M^{-1}s^{-1}$ (own assumption of slower $k_{on}$) | 0.1 $s^{-1}$ (based on *(56)*) | D=6.05*$10^{-6}$ cm²/s (based on *(55)*) | Single release event, evenly distributed over whole cell area (own assumption) | $10^6$ DA molecules (own assumption of increased DA release) |
| **Run 3** | $10^6$ $M^{-1}s^{-1}$ (based on*(56)*) | 0.01 $s^{-1}$ (own assumption of slower $k_{off}$) | D=6.05*$10^{-6}$ cm²/s (based on*(55)*) | Release from small vesicles at a random position on the edge of the cell, 30 release events every 2 s (own assumption, singular release events from different vesicles given e.g. in*(58)*) | 30000 DA molecules per release event (as in *(58)*) |
| **Run 4** | $10^4$ $M^{-1}s^{-1}$ (own assumption of slower $k_{on}$) | 0.01 $s^{-1}$ (own assumption of slower $k_{off}$) | D=6.05*$10^{-6}$ cm²/s (based on*(55)*) | Single release event, evenly distributed over whole cell area (own assumption) | $10^6$ DA molecules (own assumption of increased DA release) |
| **Run 5** | $10^4$ $M^{-1}s^{-1}$ (own assumption of slower $k_{on}$) | 0.1 $s^{-1}$ (based on*(56)*) | D=6.05*$10^{-9}$ cm²/s (own assumption of drastically reduced diffusion speed due to space below cell) | Single release event, evenly distributed over whole cell area (own assumption) | $10^6$ DA molecules (own assumption of increased DA release) |

**Table S6. Number of donors and cells (in single cell experiments)**

| Figure | | n [donors] | N [cells] |
|---|---|---|---|
| 1B | all | 6 | No single-cell data |
| 1D | DA | 12 | No single-cell data |
| | NE | 6 | |
| | E | 9 | |
| | HVA | 6 | |
| | VMA | 6 | |
| 1H | Control/5-HT | 3 | No single cell data |
| 2F | Control | 4 | 138 |
| | LPS | 4 | 49 |
| | 5-HT | 3 | 35 |
| | fMLP | 4 | 51 |
| 2I | CA | 2 | 30 |
| | FFN102 | 2 | 30 |
| 3B | Control | 3 | No single-cell data |
| | DA | 3 | |
| | 5-HT | 6 | |
| 4C | 5-HT | 5 | 246 |
| | Ketanserin | 5 | 121 |
| | CP809 | 3 | 96 |
| 4G | Control | No cells | No single-cell data, 18 ROIs |
| | Platelets | 3 | No single-cell data |
| | Platelets + Neutrophils | 3 | 44 |
| 5B | Neutrophil lysate | 5 | No single-cell data |
| | NTC | 3 | |
| 5C | DA 1 µM | 3 | No single-cell data |
| | DA 100 µM | 6 | |
| | NE 1 µM | 3 | |

| | | | |
|---|---|---|---|
| | NE 100 µM | 6 | |
| | E 1 µM | 6 | |
| | E 100 µM | 9 | |
| | 5-HT 1 µM | 5 | |
| | 5-HT 100 µM | 5 | |
| 5D | Control | 14 | No single-cell data |
| | all others | 8 | |
| 5E | Control | 13 | No single-cell data |
| | all others | 8 | |
| S2 | Neutrophils/Inhibited Neutrophils | 5 | 182/132 |
| | HEK/Inhibited HEK | 4 | 80/6 |
| | HeLa/Inhibited HeLa | 3 | 50/18 |
| S3C | 5-HT | 1 | 10 |
| | Control | 1 | 3 |
| S5 | 1 | 5 | 15 |
| | 2 | 5 | 27 |
| | 3 | 5 | 150 |
| | 4 | 5 | 16 |
| | 5 | 5 | 25 |
| S8 | Platelets + inhibited Neutrophils | 1 | 18 |
| | Platelets + Neutrophils | 1 | 16 |
| S9C | Ionomycin | 1 | 11 |
| S10 | all | 5 | No single-cell data |
